# A salt bridge-mediated resistance mechanism to FtsZ inhibitor PC190723 revealed by a cell-based screen

**DOI:** 10.1101/2022.04.06.487355

**Authors:** Ajay Kumar Sharma, Sakshi Mahesh Poddar, Joyeeta Chakraborty, Bhagyashri Soumya Nayak, Srilakshmi Kalathil, Nivedita Mitra, Pananghat Gayathri, Ramanujam Srinivasan

## Abstract

Bacterial cell division proteins, especially the tubulin homolog FtsZ, have emerged as strong targets for developing new antibiotics. Here, we have utilized the fission yeast heterologous expression system to develop a cell-based assay to screen for small molecules that directly and specifically target the bacterial cell division protein FtsZ. The strategy also allows for simultaneous assessment of the toxicity of the drugs to eukaryotic yeast cells. As a proof-of-concept of the utility of this assay, we demonstrate the effect of the inhibitors sanguinarine, berberine and PC190723 on FtsZ. Though sanguinarine and berberine affect FtsZ polymerization, they exert a toxic effect on the cells. Further, using this assay system, we show that PC190723 affects *Helicobacter pylori* FtsZ function and gain new insights into the molecular determinants of resistance to PC190723. Based on sequence and structural analysis and site-specific mutations, we demonstrate that the presence of salt-bridge interactions between the central H7 helix and beta-strands S9 and S10 mediate resistance to PC190723 in FtsZ. The single-step *in vivo* cell-based assay using fission yeast enabled us to dissect the contribution of sequence-specific features of FtsZ and cell permeability effects associated with bacterial cell envelopes. Thus, our assay serves as a potent tool to rapidly identify novel compounds targeting polymeric bacterial cytoskeletal proteins like FtsZ to understand how they alter polymerization dynamics and address resistance determinants in targets.

## Introduction

The rapid emergence and spread of antibiotic resistance to almost every antibiotic currently in use against bacteria, has resulted in an urgent need to develop new classes of antibiotics. Among the new targets with a novel mechanism of action to overcome the emerging resistance problem (Butler and Paterson 2020), one of the most potential target candidates is FtsZ, an essential bacterial cell division machinery protein (Silber et al. 2020; Wang et al. 2003; Schaffner-Barbero et al. 2012; Carro 2019; Zorrilla et al. 2021; Tripathy and Sahu 2019; Andreu et al. 2022). FtsZ is a prokaryotic homolog of tubulin, which is highly conserved and present in all bacteria and many archaea (Addinall and Holland 2002; Vaughan et al. 2004; Erickson et al. 2010; Erickson 1995; Margolin 2005; Nogales et al. 1998). FtsZ monomers self-assemble into polymeric structures in presence of GTP to give rise to FtsZ protofilaments *in vitro* and assemble into the so-called Z-ring at the division site *in vivo*. The Z-ring is the first structure to be assembled at the division site and is responsible for the recruitment of more than a dozen other accessory proteins, which comprise the divisome, the division apparatus of the cell (Bi and Lutkenhaus 1991; Ma et al. 1996; Weiss et al. 1999; Weiss 2004; Huang et al. 2013; RayChaudhuri and Park 1992; Bramhill and Thompson 1994). The Z-ring is a highly dynamic structure with a half-life of less than 10 seconds due to the constant exchange of FtsZ molecules between the ring and the soluble monomers (Anderson et al. 2004). Since the Z-ring lies at the core of the cell division process, its functional defects result in bacterial growth inhibition and cell death (Vollmer 2006; Zorrilla et al. 2021; Mahone and Goley 2020; Barrows and Goley 2021; Addinall and Lutkenhaus 1996).

FtsZ has three important protein pockets that bind small molecules and are collectively known as druggable regions. These regions are the nucleotide-binding domain (NBD), the inter-domain cleft (IDC) present between the central core helix (H7 helix) and the C-terminal domain (CTD) and the T7 loop (Löwe and Amos 1998; Kusuma et al. 2019; Andreu et al. 2022). The T7 loop of one subunit is inserted into the NBD of another subunit of FtsZ to form a catalytic site for GTP hydrolysis. Several compounds have been evaluated for their activity against FtsZ from both Gram-positive bacteria and Gram-negative bacteria. Although many exhibited only weak activity *in vivo* against Gram-negative bacteria, derivatives could be promising. These include benzamides (Haydon et al. 2008; Adams et al. 2011; Straniero et al. 2017, 2020a), trisubstituted benzimidazoles (Kumar et al. 2011), 4-bromo-1*H*-indazole derivatives (Wang et al. 2015), cinnamaldehyde and its derivatives (Domadia et al. 2007; Li et al. 2015), curcumin (Rai et al. 2008), heterocyclic molecules like guanidinomethyl biaryl compounds (Kaul et al. 2012), pyrimidine-quinuclidine scaffolds (Chan et al. 2013), 3-phenyl substituted 6,7-dimethoxyisoquinoline (Kelley et al. 2012), thiazole orange derivatives (Sun et al. 2017), viriditoxin (Wang et al. 2003), N-heterocycles such as zantrins and derivatives (Margalit et al. 2004; Nepomuceno et al. 2015).

Berberine and sanguinarine are plant-derived benzylisoquinoline alkaloids, with antibacterial activity (Wolff and Knipling 1993; Pierpaoli et al. 2021; Cernáková and Kostálová 2002; Chu et al. 2016). Berberine binds to FtsZ at a 1:1 ratio *in vitro* and exhibits dose-dependent inhibition of FtsZ assembly kinetics with a reduction in GTPase activity. Berberine is believed to target the GTP-binding pocket in the FtsZ structure and further affects the FtsZ polymer stability and Z-ring formation *in vivo* (Boberek et al. 2010; Domadia et al. 2008; Sun et al. 2014; Pradhan et al. 2021). Sanguinarine, also thought to bind the nucleotide-binding domain (NBD) in FtsZ, prevents the bundling of FtsZ protofilaments *in vitro* and perturbs the Z-ring assembly *in vivo* (Beuria et al. 2005). The precise mechanism of inhibition or the binding site of these inhibitors remains unclear.

The most potent and promising among the FtsZ inhibitors is PC190723, a 2,6-difluorinated 3-methoxybenzamide derivative, as it has proven to be efficacious in *in vivo* model of mice infected with a lethal dose of *Staphylococcus aureus* (Haydon et al. 2008). Crystal structures of FtsZ from *S. aureus* with PC190723 reveal binding at the inter-domain cleft between the central core helix (H7 helix) and the C-terminal domain (CTD) of FtsZ (Matsui et al. 2012a; Tan et al. 2012) and is the only inhibitor whose target site in FtsZ is clearly established. PC190723 binds preferentially to the T state of *S. aureus* FtsZ in which the cleft is opened, thus stabilizing the high-affinity conformation required for FtsZ polymer elongation (Matsui et al. 2012a; Fujita et al. 2017). Thereby, PC190723 reduces the critical concentration for polymerization and induces FtsZ assembly (Andreu et al. 2010; Elsen et al. 2012). Although the resistance of streptococci, enterococci and many Gram-negative bacteria to PC190723 (Haydon et al. 2008; Kaul et al. 2013b) and poor drug and pharmacokinetic properties have hampered the pre-clinical trials with PC190723 (Stokes et al. 2013; Kaul et al. 2013a), phase-I tests are in progress by TAXIS pharmaceuticals (Monmouth Junction, NJ, USA) for the benzamide class of drug TXA709 (Butler and Paterson 2020; TAXIS Pharmaceuticals 2020).

Thus, despite several years of intense research on FtsZ inhibitors, a commercially viable antibiotic that targets FtsZ is not yet available. One of the major hindrances has been the validation that FtsZ is the actual direct target of these small molecule compounds (Kusuma et al. 2019; Zorrilla et al. 2021; Schaffner-Barbero et al. 2012). Although several other small molecules have been shown to inhibit bacterial cell division, presumably by directly inhibiting FtsZ function, many of them have also been shown to be due to indirect effects on membranes (Hurley et al. 2016). Several methods have been used to ascertain FtsZ as the target of the drug, and the various approaches have been reviewed in detail by many (Kusuma et al. 2019; Silber et al. 2020; Zorrilla et al. 2021; Andreu et al. 2022). Andreu et al. (2022) have recently proposed a streamlined experimental protocol for the screening and characterization of FtsZ inhibitors. A scheme of methods to be followed, from the identification of hit compounds to the development of a lead compound, has also been proposed. These include several assays for monitoring effects on FtsZ polymerization, cytological effects on eukaryotic cells, pharmacological properties and structural optimization (Andreu et al. 2022; Schaffner-Barbero et al. 2012). Any small molecule antagonist identified thus should have no off-target effects and minimally affect eukaryotic cells. A single-step screening method that identifies small molecules, directly targeting FtsZ assembly and as well as selects out compounds probably toxic to eukaryotic cells would be ideal. Moreover, the screen should be easily scalable to high-throughput platforms.

We had previously established *Schizosaccharomyces pombe* as a useful model system to study bacterial cytoskeletal proteins. We had shown that A22, a small molecule inhibitor of the actin homologue, MreB, prevented its polymerization in yeast cells as well (Srinivasan et al. 2007). Further, we reported that FtsZ assembles into ring-like structures with similar characteristics to Z-rings in bacterial cells when expressed in *S. pombe* or *Saccharomyces cerevisiae* (Srinivasan et al. 2008). Moreover, the dynamics of chloroplast FtsZs have also been successfully studied using the heterologous fission yeast expression system (TerBush and Osteryoung 2012; Yoshida et al. 2016; TerBush et al. 2018). These observations prompted us to explore if the heterologous expression system using *S. pombe* could serve as a platform for testing and screening chemical compounds targeting FtsZ directly.

In this study, we show that FtsZ from two pathogenic bacterial species, *S. aureus* and *Helicobacter pylori*, expressed in fission yeast, assemble into dynamic ring-like and spiral polymers, respectively. We test the effect of known small molecule inhibitors of FtsZ, sanguinarine, berberine and PC190723 on the assembly of FtsZ polymers. Herein, we thus develop a proof-of-concept of a cell-based screening method that is easily scalable to existing high-throughput platforms. The method combines the ability to establish the specificity of the target by directly assaying the polymerization of FtsZ in the cellular milieu without a tedious protein purification step, as well as assessing toxicity to yeast cells in a single step. We find that although sanguinarine and berberine affected FtsZ polymerization, they also affected yeast cell physiology. We show that while all three inhibitors exert an effect on FtsZ assembly, PC190723 specifically promotes the assembly of FtsZ polymers of *S. aureus* and *H. pylori*. Further, we probed the determinants of resistance to PC190723. Based on FtsZ sequence and structure analysis, we show that salt-bridge interactions between the central H7 helix and S9 / S10 β-sheet prevent the inhibitor from accessing the inter-domain cleft (IDC). This appears to be a sequence-determined resistance probably prevalent in many Gram-negative bacteria. We thus demonstrate the utility of the heterologous yeast system (*S. pombe*) for identifying chemical compounds directly targeting FtsZ, evaluating them for toxicity to eukaryotic cells and studying the molecular determinants of drug resistance.

## Results

### S. aureus and H. pylori FtsZ assemble into polymeric structures in fission yeast

We expressed FtsZ from two pathogenic species, *S. aureus* and *H. pylori* (hereafter referred to as SaFtsZ for *S. aureus* FtsZ and HpFtsZ for *H. pylori* FtsZ, respectively) as C-terminal GFP fusions in *S. pombe*. Expression of FtsZ in *S. pombe* was achieved from a pREP42-GFP vector (a medium strength thiamine repressible promoter *nmt41/42* containing vector; (Basi et al. 1993; Moreno et al. 2000) by growing cultures in minimal medium in the absence of thiamine (induction medium) as reported earlier (Srinivasan et al. 2008). Epifluorescence microscopy showed that SaFtsZ assembled predominantly into spot-like structures (**Fig. 1A**) and were similar in appearance to that assembled by FtsZ from *Escherichia coli* (EcFtsZ) (**Fig. 1B**). Assembly of SaFtsZ into ring-like structures was only discernible by super-resolution microscopy using structural illumination (3D-SIM) (**Fig. 1C**), whereas ring-like structures of EcFtsZ were visualized by deconvolution of images (**Fig. 1B iii**). Interestingly, HpFtsZ assembled into linear cable-like structures as well as twisted polymers that were curled and spiral in appearance (**Fig. 1D**). The spiral filaments were more clearly visualized by deconvolution of the images (**Fig. 1D iii and 1E**). Further, super-resolution imaging using 3D-SIM clearly revealed that HpFtsZ assembles into spiral filaments in fission yeast (**Fig. 1F**). Spiral polymers appeared early, at 16 – 18 hours after induction of expression (absence of thiamine), and linear cables appeared later at 20 – 22 hours **(Fig. S1)**. The smooth linear polymers possibly arise from lateral association and bundling of FtsZ filaments (Monahan et al. 2009), but the factors determining the two forms in yeast cells remain unclear.

**Figure 1.**
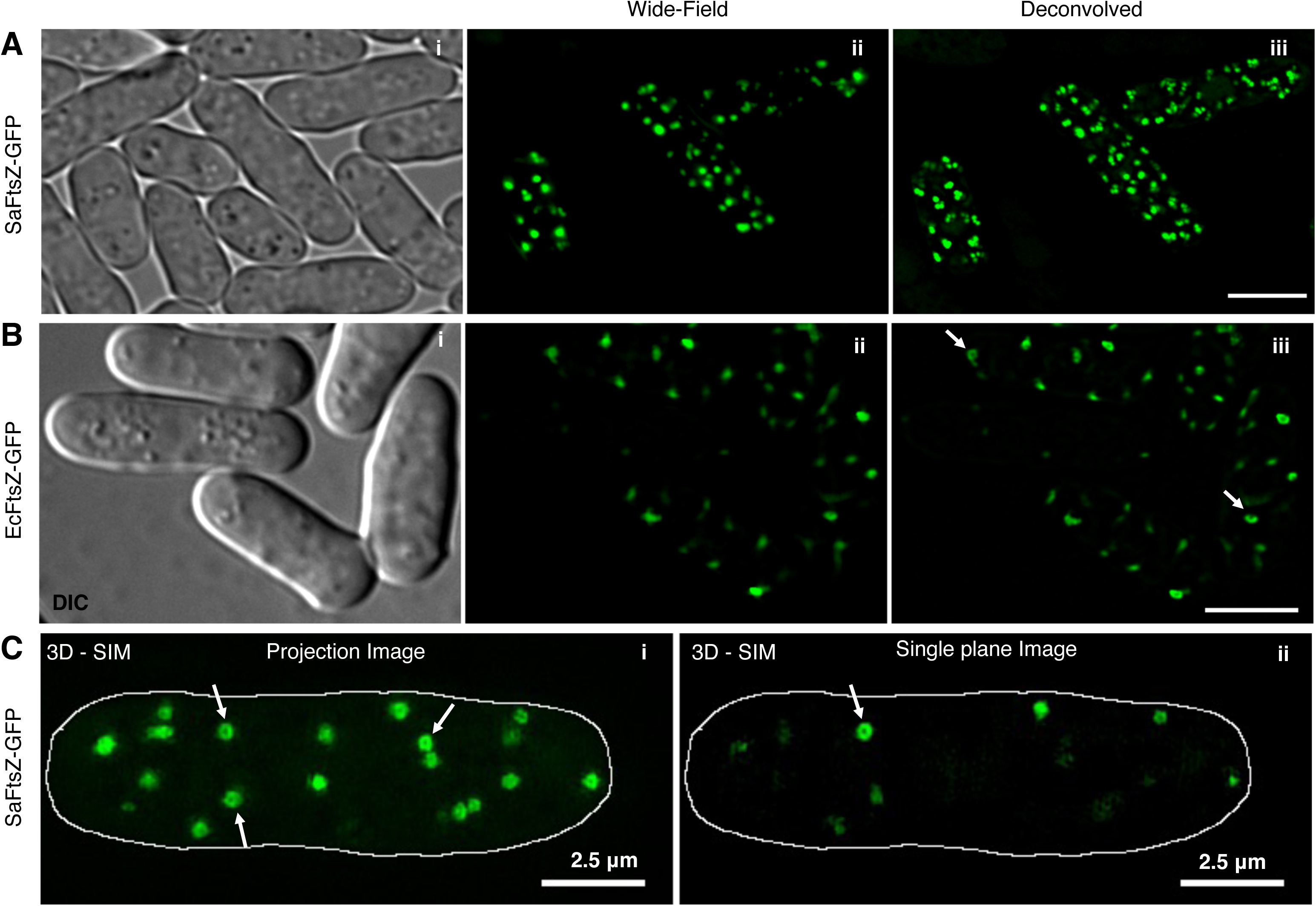
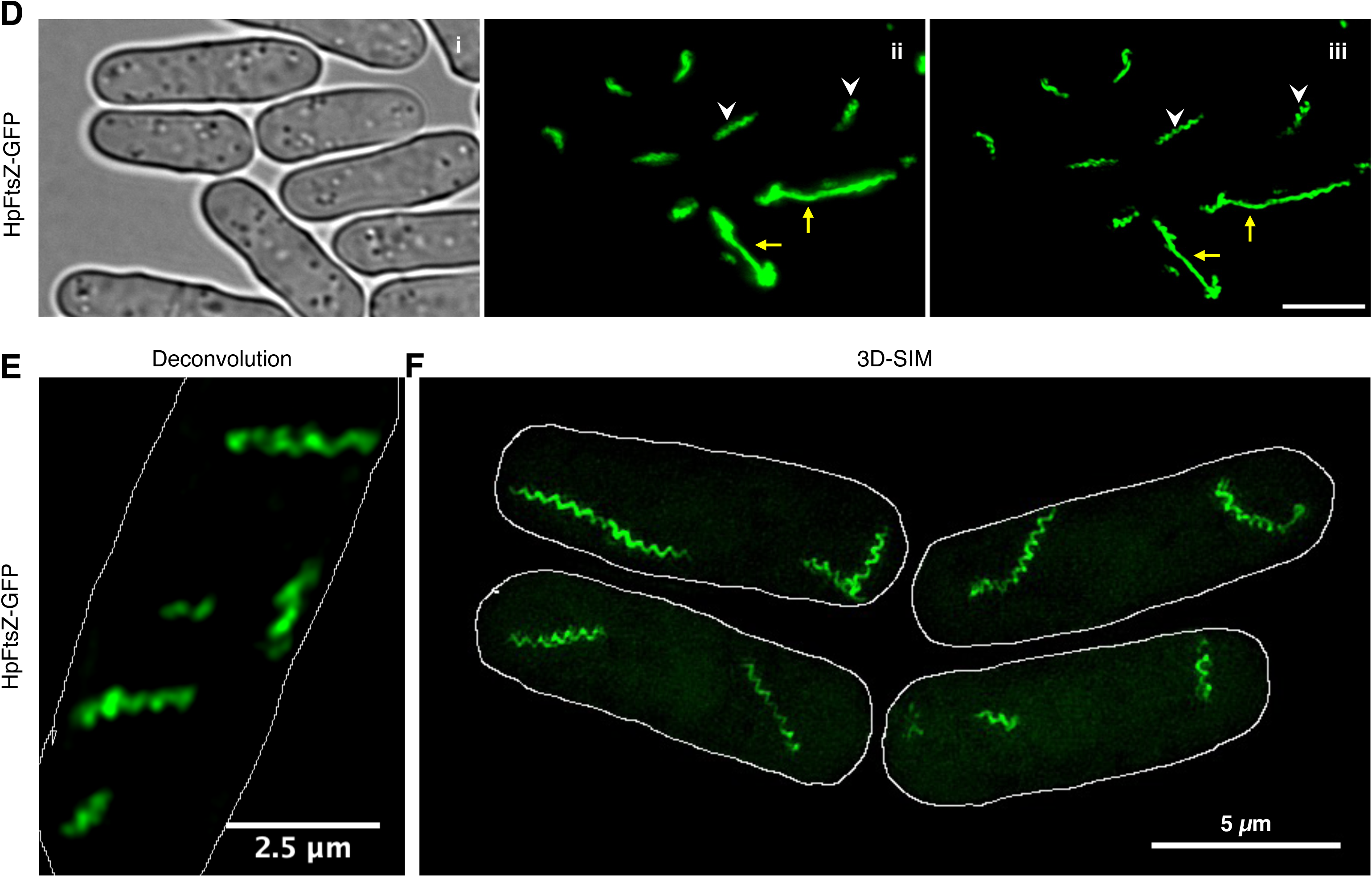
Polymerization of SaFtsZ, EcFtsZ and HpFtsZ in fission yeast. The medium-strength *nmt42* promoter was used for the expression of different bacterial FtsZ-GFP fusion constructs in *S. pombe*. *S. pombe* cultures expressing FtsZ-GFP fusions were grown for 16–20 h at 30°C in the absence of thiamine to allow the expression of the proteins, mounted onto 1.6% agarose slide, and imaged using an **(A, B, E and D)** epifluorescence microscope or **(C and F)** super-resolution using 3D-SIM. **(A)** SaFtsZ and **(B)** EcFtsZ assembles into spots and cable-like structures in fission yeast (panels ii and iii). Arrows indicate ring-like structures. **(C)** 3D-SIM images of SaFtsZ. **(C i)** Maximum intensity projection image and **(C ii)** a single plane image reveal ring-like structures of SaFtsZ. Arrows indicate visible SaFtsZ rings. **(D)** HpFtsZ forms cable-like structures. Spiral filaments of HpFtsZ are indicated by white arrowheads, and linear cables are marked with yellow arrowheads. DIC or phase-contrast images are shown in **Ai, Bi & Di**. **(C)** Deconvolved image of HpFtsZ and **(F)** 3D-SIM images of HpFtsZ show the assembly of HpFtsZ into helical filaments in fission yeast. Scale bar represents 5 μm for all images **(A, B, D and F)** except for the image in **C** and **E**, where it represents 2.5 μm.

We had previously shown that the diameter of EcFtsZ rings assembled in yeast were 0.48 ± 0.042 μm (SEM ± SD) (Srinivasan et al. 2008). Quantification of the diameter of the rings assembled by SaFtsZ revealed that they had a mean diameter of 0.29 ± 0.04 μm (SEM ± SD) and were thus smaller than EcFtsZ rings **(Fig. S2A)**. The spiral polymers of HpFtsZ had a mean diameter of 0.35 ± 0.01 μm and a pitch of 0.41 ± 0.03 μm (SEM ± SD) **(Fig. S2B i and ii)**. These results show that FtsZ from both species *S. aureus* (SaFtsZ) and *H. pylori* (HpFtsZ), when expressed in fission yeast cells, assemble into polymeric structures like many other FtsZs.

### Sanguinarine and berberine prevent FtsZ assembly in fission yeast but also affect yeast cell morphology

Sanguinarine and berberine have been shown to perturb FtsZ ring assembly in *E. coli* (Beuria et al. 2005; Boberek et al. 2010; Domadia et al. 2008; Liu et al. 2017). Although sanguinarine has been reported to inhibit septation and cell division in *S. aureus* (Obiang-Obounou et al. 2011), its effects on SaFtsZ or HpFtsZ have not yet been reported. We, therefore, tested if any of these molecules exhibited any direct effects on FtsZ expressed in yeast cells. Fission yeast cultures were grown at 30°C in the absence of thiamine for 10 – 12 hours to induce the expression of the bacterial FtsZ-GFP fusions. DMSO or drugs were subsequently added to the cultures and further grown at 30°C for another 10 – 12 hours prior to imaging.

A microscopic examination of fission yeast cells expressing SaFtsZ (**Fig. 2A**), EcFtsZ (**Fig. 2B**) or HpFtsZ (**Fig. 2C**) in the presence of sanguinarine (**Fig. 2 A ii – C ii**) or berberine (**Fig. 2 A iii – C iii**) revealed that assembly of FtsZ into polymers was inhibited in the presence of the drugs as compared to untreated (DMSO added) cells (**Fig. 2 A i – C i**). A significant reduction in the percentage of cells that exhibited FtsZ assemblies (spots/ dots in the case of SaFtsZ or EcFtsZ and filaments in the case of HpFtsZ) in the presence of sanguinarine or berberine as compared to control untreated cells (**Fig. 2 D i – iii and F i – iii**) was observed (≥ 500 cells counted for each replicate; N = 3). To obtain a quantitative measure of the effect of drugs on FtsZ assembly, we compared the total number of spots per cell in the case of SaFtsZ and EcFtsZ from treated and untreated cells. For statistical analysis, we used both estimation statistics (effect size and 95 % CI) and significance testing by null hypothesis statistical tests (Claridge-Chang and Assam 2016). When treated with sanguinarine, a reduction in the number of spots per cell was observed in both SaFtsZ and EcFtsZ, with an effect size of −4.82 [95% CI −5.7, −3.9] and −5.4 [95% CI −9.7, −1.1] respectively (**Fig. 2E i and ii**). Similarly, the addition of berberine also exhibited a similar reduction in the number of spots per cell in both SaFtsZ and EcFtsZ, with an effect size of −5.6 [95% CI −7.9, −3.3] and −12.4 [95% CI −19.5, −5.4] respectively (**Fig. 2G i and ii**). Since HpFtsZ assembled into filaments, we compared the total length of the polymer per cell in the presence or absence of the drugs. Both sanguinarine and berberine resulted in a significant reduction of total polymer length per cell with an effect size of −45.3 [95% CI −60, −30] and −92 [95% CI −151, −33.5], respectively (**Fig. 2 E iii and G iii**).

**Figure 2.**
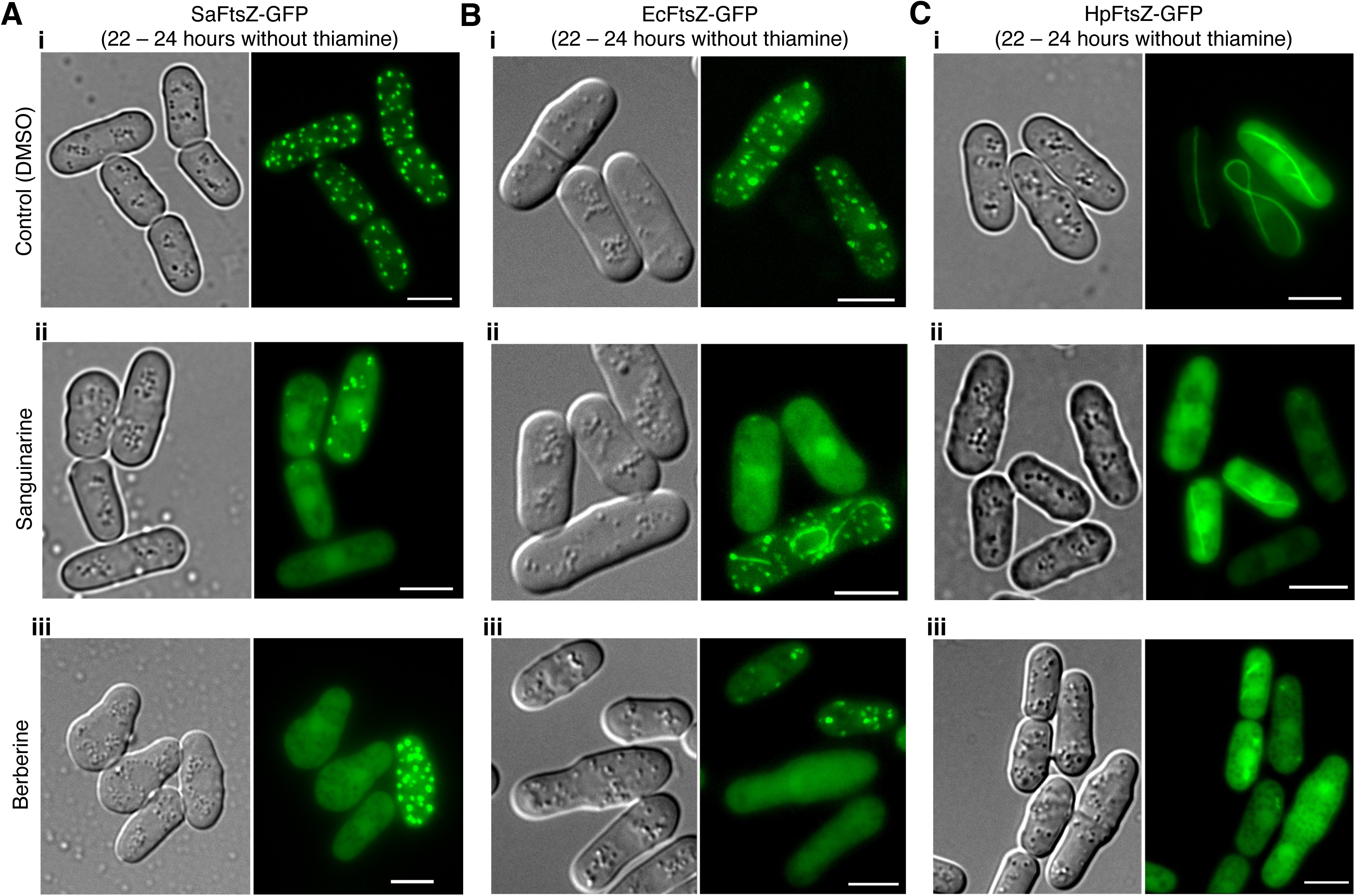
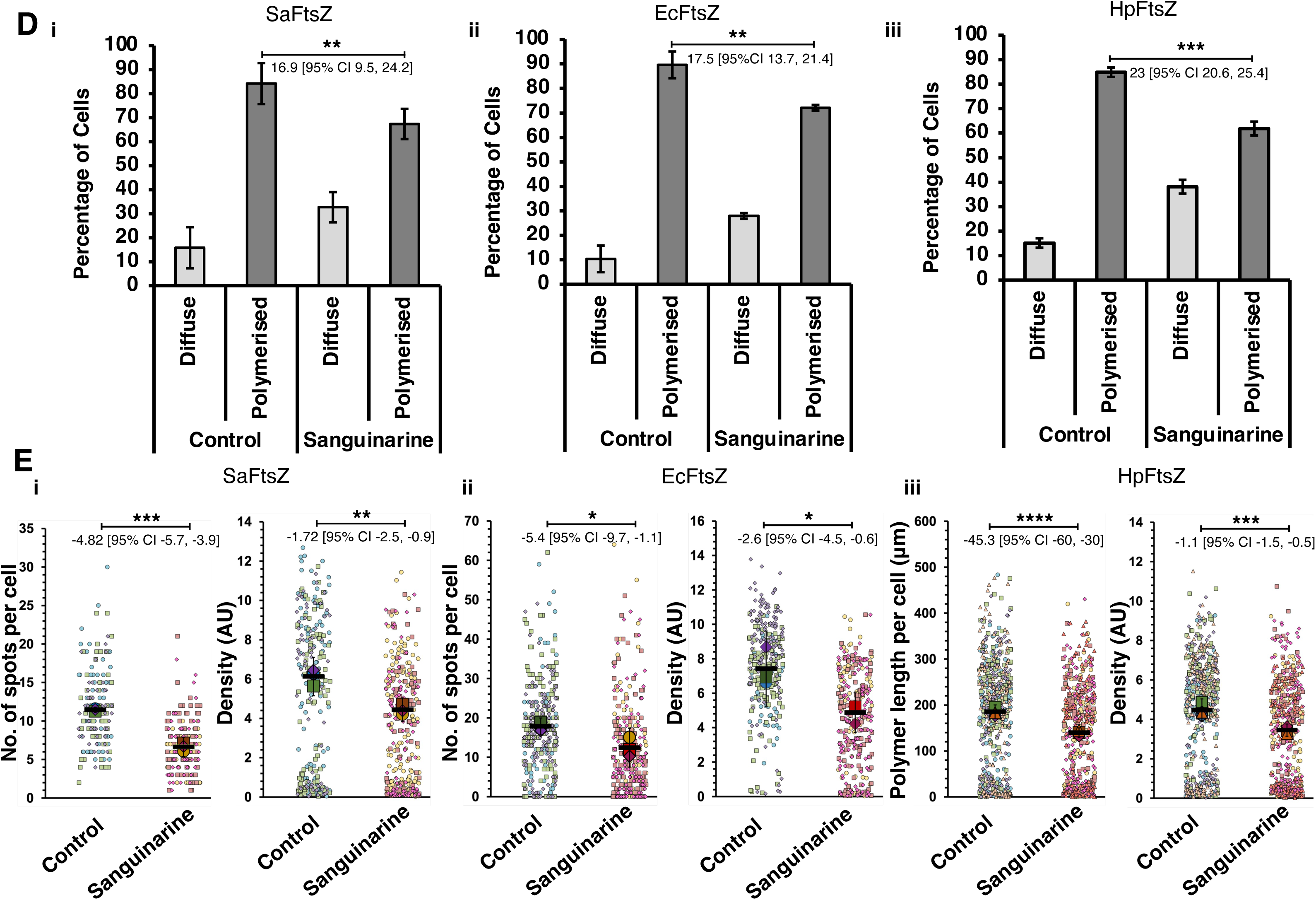
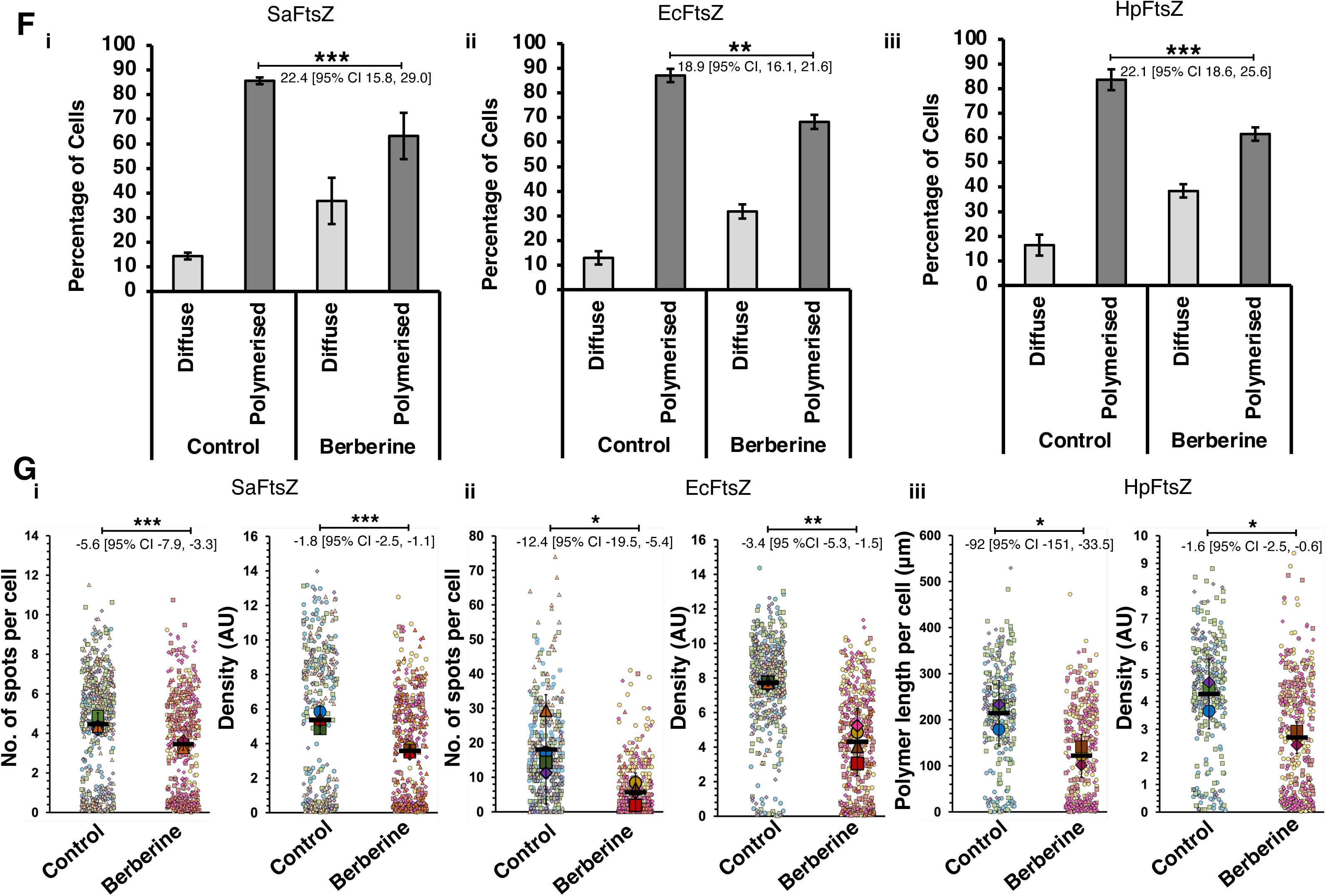
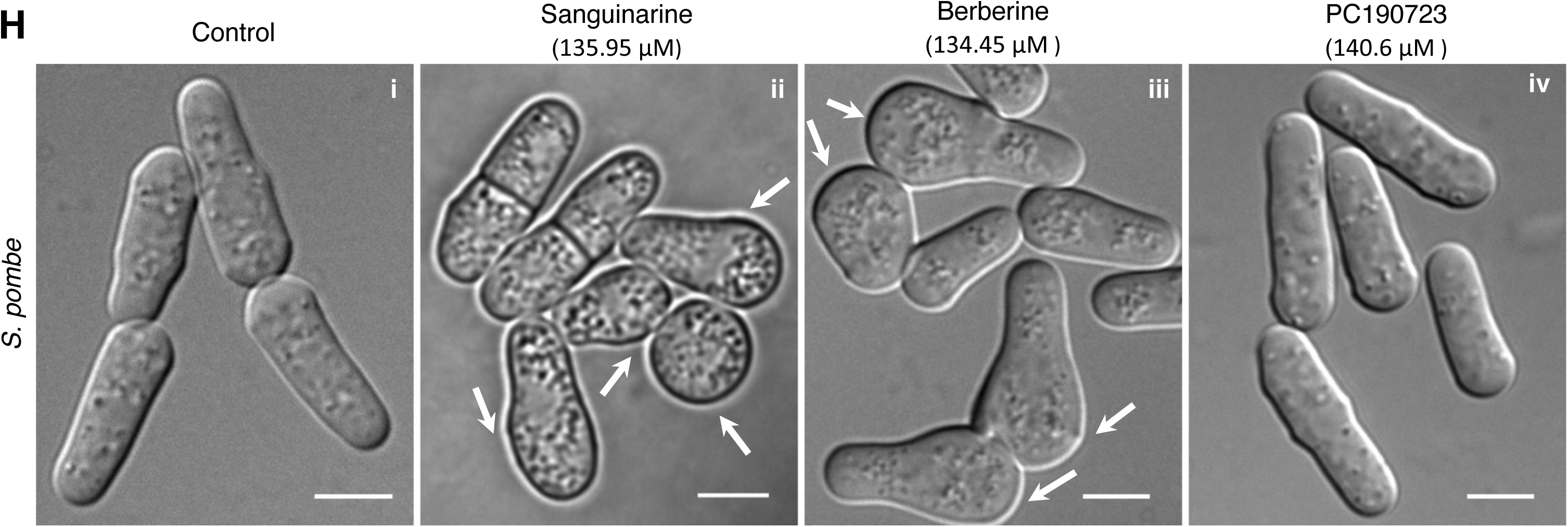
Inhibition of FtsZ polymerisation by sanguinarine and berberine. Fission yeast cultures expressing different bacterial FtsZ-GFP fusions were grown at 30°C in the absence of thiamine for 10 – 12 hours before DMSO or drugs were added to the cultures. The cultures were further grown at 30°C for another 10 – 12 hours, either in the absence or presence of the drug before imaging or counting. Cells were mounted onto 1.6% agarose slides, and an epifluorescence microscope was used for the imaging. In control (untreated) cultures, an equivalent amount of DMSO was added. **(A i – C i)** Control (DMSO-treated) cells showed normal assembly of FtsZ polymers in fission yeast. **(A i)** SaFtsZ **(B i)** EcFtsZ and **(C i)** HpFtsZ. **(A ii – C ii)** Sanguinarine (20 μM) treated cells showed a significant reduction in **(A ii)** SaFtsZ, **(B ii)** EcFtsZ and **(C ii)** HpFtsZ. **(A iii – C iii)** Berberine (53.79 μM) inhibited the polymerization of **(A iii)** SaFtsZ, **(B iii)** EcFtsZ and **(C iii)** HpFtsZ. Scale bar represents 5 μm. **(D – G)** Quantification of the effects of sanguinarine **(D and E)** and berberine **(F and G)** on polymerization of SaFtsZ **(i)**, EcFtsZ **(ii)** and HpFtsZ **(iii)**, respectively. **(D and F)** show the percentage of cells having polymers in the presence or absence of sanguinarine **(D)** or berberine **(F)**. Light gray bars and dark gray bars represent cells having diffuse fluorescence and cells containing FtsZ polymers, respectively. **(E and G)** Superplots showing the quantification of the number of spots per cell and density, which is a measure for the amount of assembled cytoskeleton per unit area in a given cell for **(i)** SaFtsZ or **(ii)** EcFtsZ or total polymer length per cell for and density for **(iii)** HpFtsZ when treated with **(E)** sanguinarine or **(G)** berberine. For the quantification of spots in **(E)**, the number of cells counted for each replicate for SaFtsZ and EcFtsZ were ≥ 44 and ≤ 72; N = 3 and ≥ 64 and ≤ 130; N = 3, respectively. In the case of HpFtsZ, polymer length per cell was measured from ≥ 112 and ≤ 147 cells for each replicate (N = 4). For density measurement, the number of cells counted for each replicate for SaFtsZ, EcFtsZ, and HpFtsZ was ≥ 68 and ≤ 167; N = 3, ≥ 64 and ≤ 130; N = 3 and ≥ 112 and ≤ 147; N = 4 respectively. For the quantification of spots in **(G)**, the number of cells counted for each replicate for SaFtsZ and EcFtsZ were ≥ 42 and ≤ 82; N = 4 and ≥ 48 and ≤ 136; N = 4, respectively. In the case of HpFtsZ, polymer length per cell was measured from ≥ 91 and ≤ 182 cells for each replicate (N = 3). For density measurement, the number of cells counted for each replicate for SaFtsZ, EcFtsZ, and HpFtsZ was ≥ 75 and ≤ 151; N = 4, ≥ 48 and ≤ 136; N = 4 and ≥ 91 and ≤ 182; N = 3 respectively. The black bars represent the mean values derived from independent biological replicates, the large markers represent the mean values of each replicate, and the small markers represent individual cells. The error bars shown are inferential and represent Mean ± 95% CI. Asterix ***, ** and * indicate P-values ≤ 0.001, ≤ 0.01 and < 0.05 respectively. P-values were calculated using the two-sided proportions z-test **(D and F)** or a two-sided unpaired Student’s t-test **(E and G)**. The difference in the means between the untreated and treated populations is also indicated as effect size [95% CI, lower bound, upper bound]. Biological replicates are independent liquid cultures starting from a fresh patch of yeast cells grown from frozen glycerol stocks. **(H)** Berberine and sanguinarine, but not PC190723, affect yeast cell morphology at high concentrations. *S. pombe* cultures were grown **(i)** in the absence of the drug or in the presence of **(ii)** sanguinarine (135.95 μM) or **(iii)** berberine (134.45 μM) or **(iv)** PC190723 (140.6 μM) and mounted onto 1.6% agarose slides and imaged. In control (untreated) cultures, an equivalent amount of DMSO was added. Fission yeast cells formed bulges at the poles, as indicated by arrowheads. Scale bar represents 5 μm.

In addition, we also quantified density, which is a measure of assembled cytoskeleton per unit area in a given cell (Henty-Ridilla et al. 2014; Higaki 2017). We had recently used density as a measure to successfully quantify assembly defects of an ATP hydrolysis mutant of *Spiroplasma citri* MreB (Pande et al. 2022). Cells expressing SaFtsZ, EcFtsZ or HpFtsZ and treated with the drugs had lower density values as compared to control untreated cells, suggesting an inhibitory effect of the drugs on cytoskeleton assembly. In the case of sanguinarine, the effect sizes were −1.72 [95% CI −2.5, −0.9], −2.6 [95% CI −4.5, −0.6] and −1.1 [95% CI −1.5, −0.5] for SaFtsZ, EcFtsZ and HpFtsZ respectively (**Fig. 2E i – iii**). The effect size in the case of berberine-treated cells were −1.8 [95% CI −2.5, −1.1], −3.4 [95 %CI −5.3, −1.5] and −1.6 [95% CI −2.5, −0.6] for SaFtsZ, EcFtsZ and HpFtsZ respectively (**Fig. 2G i – iii**). These results suggest that sanguinarine and berberine had an effect on the assembly of FtsZ from all three species, *S. aureus*, *H. pylori* and *E. coli* when expressed in fission yeast cells.

Although we did not observe any growth defect in yeast cells at lower concentrations of the drugs, earlier studies have suggested that yeast cells possibly require higher concentrations of drugs than used for mammalian cells due to the presence of the cell wall, which is particularly thick in *S. pombe* (Benko et al. 2017; Pérez and Ribas 2004). We thus explored the possibility of cell toxicity to yeast cells at higher concentrations of the drugs. At higher concentrations, both sanguinarine (135.95 μM) and berberine (134.45 μM) induced significant morphological defects in yeast cells (**Fig. 2H**). However, even at higher concentrations, neither of the drugs showed any visible effect on yeast microtubules (**Fig. S3 A and B**).

Further, in order to test if the effects of sanguinarine and berberine were specific to FtsZ polymerization, we tested its activity on the *E. coli* actin homologue MreB. Sanguinarine and berberine showed no effects on MreB assembly **(Fig. S3C iii and iv)**. However, A22, a known inhibitor of MreB (Gitai et al. 2005; Iwai et al. 2002), as previously reported (Srinivasan et al. 2007), completely inhibited the polymerization of EcMreB in fission yeast **(Fig. S3C ii)**.

### PC190723 induces polymerization of both SaFtsZ and HpFtsZ, but not EcFtsZ

Several studies have shown that PC190723 binds SaFtsZ, and crystal structures have revealed the binding site to be the inter-domain cleft of FtsZ (Haydon et al. 2008; Matsui et al. 2012a; Tan et al. 2012). Further, PC190723 preferentially binds to the T state of SaFtsZ, inhibits the GTPase activity of FtsZ, induces polymerization and stabilizes FtsZ filaments (Andreu et al. 2010; Elsen et al. 2012; Miguel et al. 2015; Fujita et al. 2017). We, therefore, decided to test PC190723 for its effects on SaFtsZ, HpFtsZ and EcFtsZ expressed in fission yeast cells. The effect of PC190723 was tested at various concentrations (ranging from 14.06 μM to 70.27 μM) on the assembly of FtsZ polymers. At 8 – 10 hours after growth in the induction medium (EMM without thiamine), the majority of cells expressing SaFtsZ or HpFtsZ showed only diffuse FtsZ molecules in the cytoplasm and had not yet assembled into structures. Therefore, *S. pombe* cells carrying SaFtsZ or HpFtsZ constructs were grown at 30°C in the absence of thiamine for 8 – 10 hours before DMSO or PC190723 was added to the cultures. PC190723 or DMSO was added to cultures at this point and incubated at 30°C for further 4 hours. In cultures where DMSO was added, 446 ± 5 (SEM; 500 cells counted for each replicate; N = 3) cells expressing SaFtsZ (**Fig. 3 A and B ii**) and 378 ± 4 (SEM; 500 cells counted for each replicate; N = 3) cells expressing HpFtsZ (**Fig. 3 D and E ii**) were found to have only diffuse FtsZ molecules in the cytoplasm.

**Figure 3.**
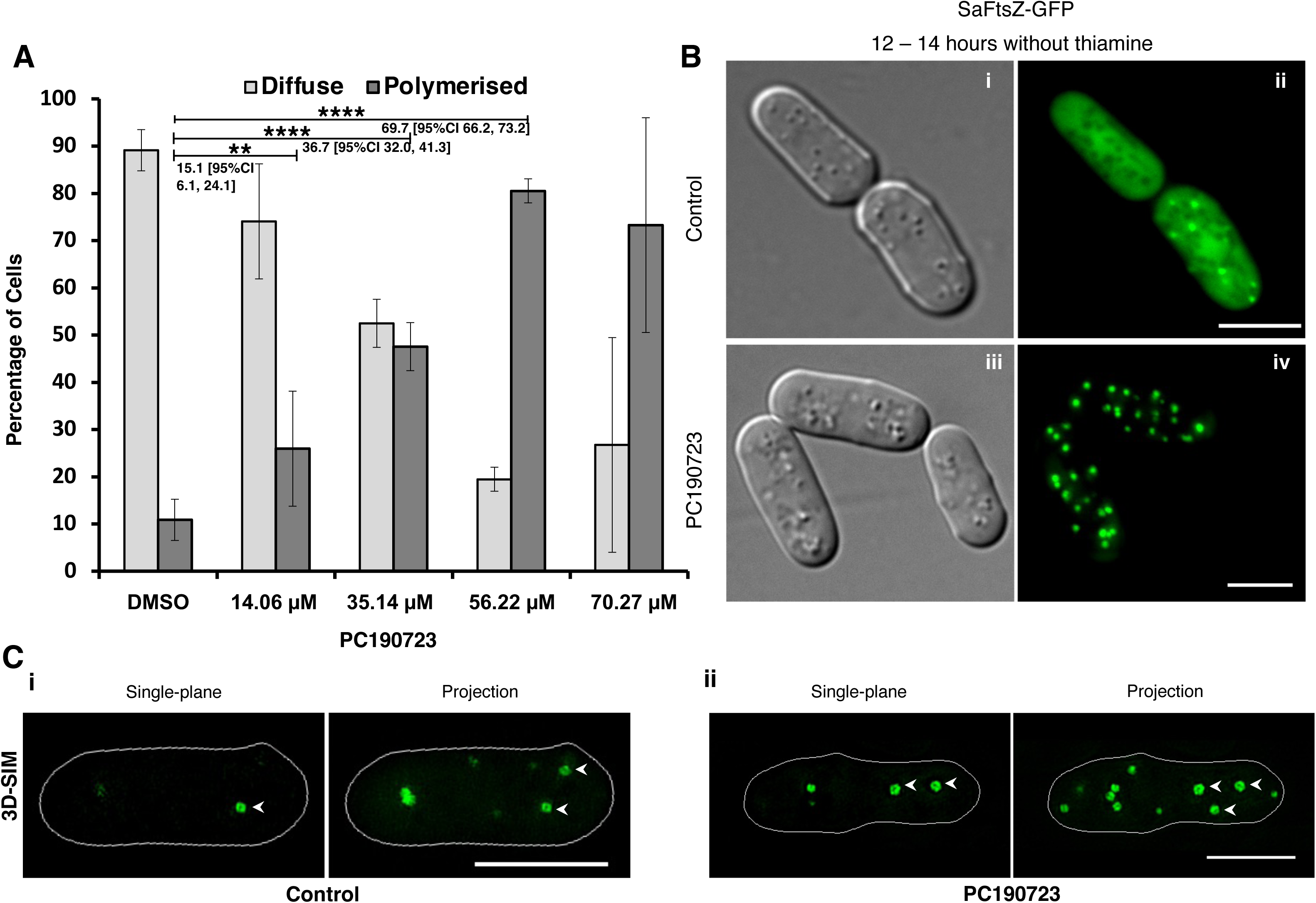
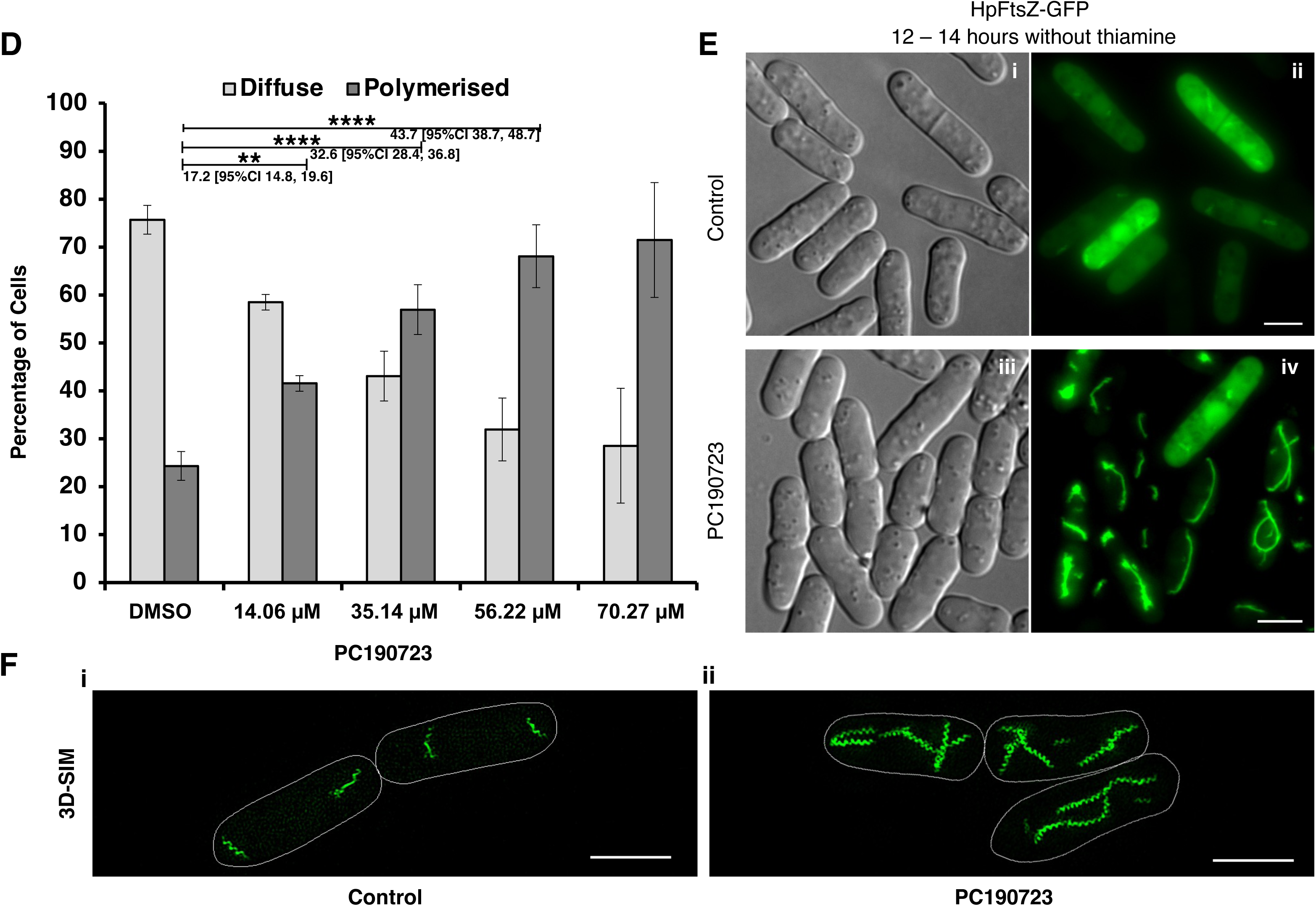
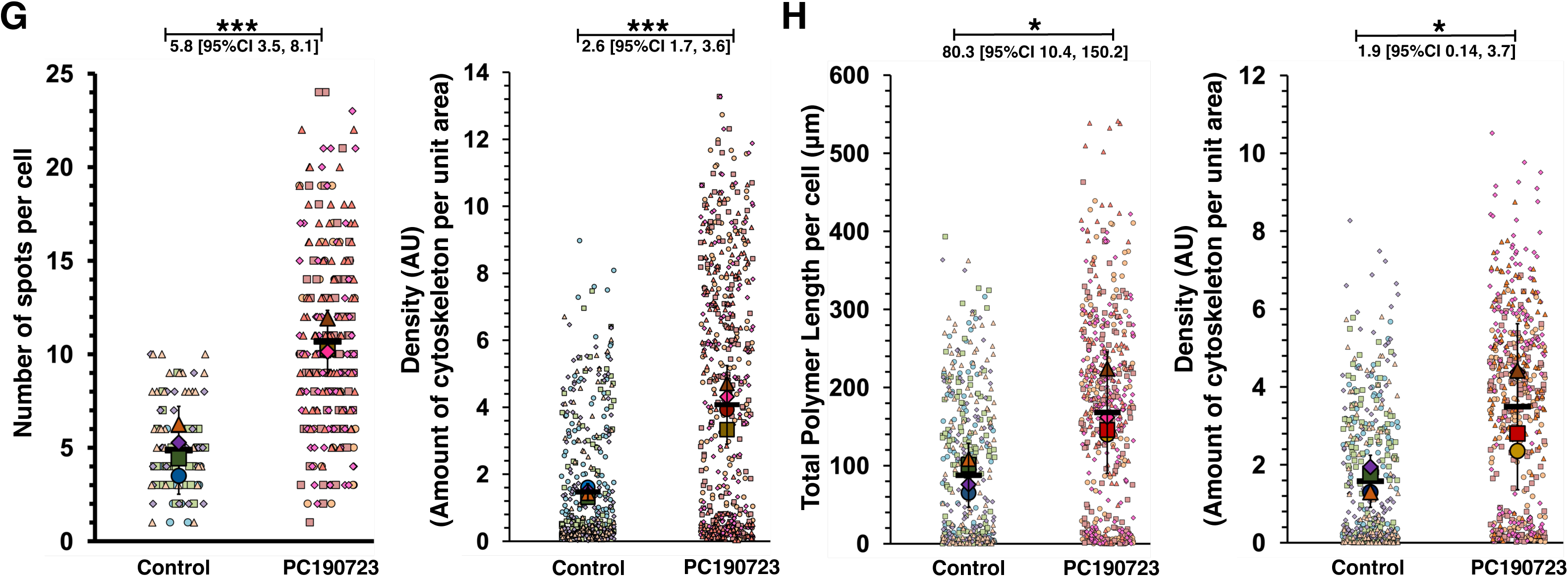
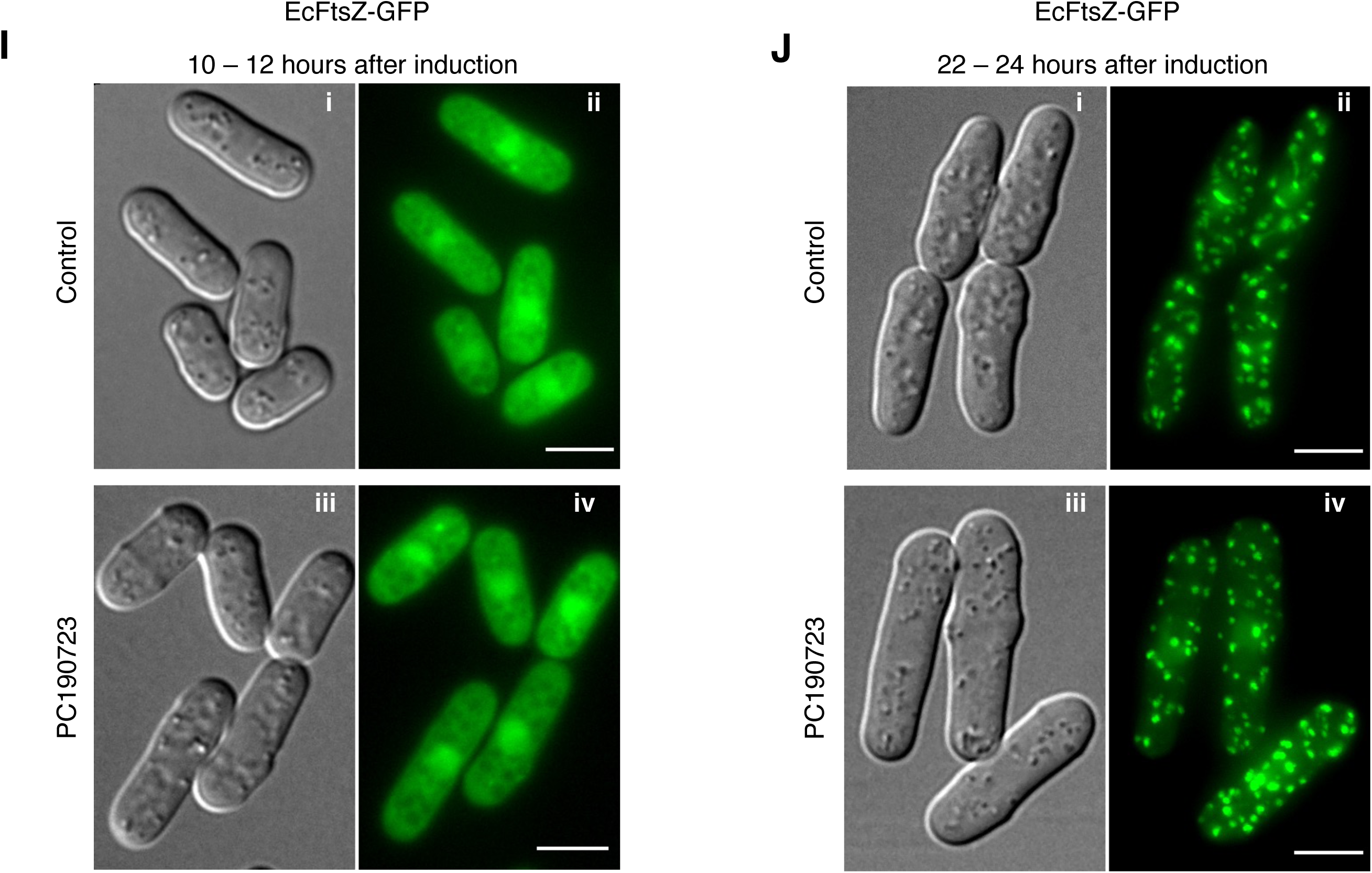
PC190723 induces polymerization of SaFtsZ and HpFtsZ in fission yeast. *S. pombe* cells expressing SaFtsZ, HpFtsZ or EcFtsZ were grown at 30°C in the absence of thiamine for 8 – 10 hours before DMSO or PC190723 was added to the cultures. The cultures were further incubated at 30°C for 4 hours before imaging or counting. **(A)** Dose-dependent effect of PC190723 on polymerization of SaFtsZ. Graph showing percentage of cells having diffuse (light gray) and polymerized (dark gray) SaFtsZ in fission yeast in the absence (DMSO) or in the presence of PC190723 ranging from 14.06 – 70.27 μM concentration. **(B)** Images showing the effect of PC190723 at 56.2 μM on SaFtsZ. At a time-point when control (DMSO treated) cells expressing SaFtsZ-GFP show diffuse fluorescence (panel ii), cells grown with PC190723 (56.2 μM) result in the early appearance of SaFtsZ assemblies (panel iv). **(C)** 3D-SIM images of SaFtsZ showing ring-like structures in both the absence **(i)** and presence **(ii)** of PC190723. **(D)** Dose-dependent effect of PC190723 on polymerization of HpFtsZ. Graph showing the percentage of cells having diffuse (light gray) and polymerized (dark gray) HpFtsZ in fission yeast in a range of PC190723 (14.06 – 70.27 μM) concentrations. **(E)** Images showing the effect of PC190723 at 56.2 μM on HpFtsZ. At a time-point when control (DMSO treated) cells expressing HpFtsZ-GFP show diffuse fluorescence (panel ii), cells grown with PC190723 (56.2 μM) result in the early appearance of HpFtsZ polymers (panel iv). **(F)** 3D-SIM images of HpFtsZ polymers showing spiral structures in both the absence **(i)** or presence **(ii)** of PC190723. **(G)** Superplots showing the quantification of the number of spots per cell and density (SaFtsZ), which is a measure for the amount of assembled cytoskeleton in a given cell in the presence or absence of PC190723. For the quantification of spots, the number of cells counted for each replicate was ≥ 18 and ≤ 85; N = 4, and for density measurement, the number of cells counted for each replicate was ≥ 86 and ≤ 176; N = 4. **(H)** Superplots for HpFtsZ showing the quantification and the effect of PC190723 on total polymer length per cell and density, a measure for the amount of assembled cytoskeleton per unit area in a given cell. For the quantification of polymer length and density measurement, the number of cells counted for each replicate was ≥ 104 and ≤ 159; N = 4. **(I) and (J)** Images showing the effect of PC190723 at 56.2 μM on EcFtsZ. **(I)** Unlike Sa or HpFtsZ, cells expressing EcFtsZ-GFP and grown in the presence of PC190723 (56.2 μM) exhibit diffuse fluorescence similar to that of control (DMSO treated) cells at early time points. *S. pombe* cells expressing EcFtsZ were grown at 30°C in the absence of thiamine for 4 – 6 hours before DMSO or PC190723 was added to the cultures. The cultures were further incubated for 4 hours at 30°C before imaging or counting. **(J)** At longer induction times (20 hours), both control (DMSO) and PC190723 treated cells formed EcFtsZ structures. DMSO or PC190723 was added to the cultures after 6 - 8 hours of growth in the absence of thiamine and incubated further at 30°C. The black bars represent the mean values derived from independent biological replicates, the large markers represent the mean values of each replicate, and the small markers represent individual cells. The error bars shown in **A** and **D** are inferential and represent the 95% CI. ****, ***, ** and * indicate P-values ≤ 0.0001, ≤ 0.001, ≤ 0.01 and < 0.05 respectively. P-values were calculated using the two-sided proportions z-test in **A** and **D**. A two-sided unpaired Student’s t-test was used to calculate P-values in **G** and **H**. The difference in the means between the untreated and treated populations is also indicated as effect size [95% CI, lower bound, upper bound]. Scale bar represents 5 μm.

On the contrary, cultures treated with PC190723 (35 μM or more) had a significantly more proportion of cells with FtsZ assembled into polymers as compared to untreated cells (DMSO) for SaFtsZ (**Fig. 3 A and B iv**) and HpFtsZ (**Fig. 3 D and E iv**). PC190723 did not alter the assembly of SaFtsZ into ring-like structures or HpFtsZ into spiral filaments (**Fig. 3 C and F; Fig. S2**). Further, with an increasing concentration of PC190723, there was a concomitant increase in the proportion of cells that contained polymerized FtsZ in the case of SaFtsZ (**Fig. 3A**) and HpFtsZ (**Fig. 3C**). Quantification of the number of spots per cell showed an increase in the number of SaFtsZ spots in the presence of PC190723 with an effect size of 5.8 [95%CI 3.5, 8.1] (cells counted for each replicate was ≥ 18 and ≤ 85; N = 4). Moreover, the amount of assembled cytoskeleton per unit area in a cell also increased, with an effect size of 2.6 [95%CI 1.7, 3.6] (cells counted for each replicate was ≥ 86 and ≤ 176; N = 4), when treated with PC190723 (**Fig. 3G**). Similarly, in the case of ΗpFtsZ, the total length of the polymer per cell and density were higher in the presence of PC190723. The effect size for polymer length and density were 80.3 [95%CI 10.4, 150.2] and 1.9 [95%CI 0.14, 3.7] (cells counted for each replicate was ≥ 104 and ≤ 159; N = 4), respectively (**Fig. 3H**). However, no such difference was observed in the absence or presence of PC190723 in cells expressing EcFtsZ (**Fig. 3 I and J**) and is consistent with a recent study that concluded that the inhibitory activity of PC190723 on *E. coli* was independent of FtsZ (Khare et al. 2019). These observations suggested that PC190723 resulted in an early assembly of FtsZ into polymers in the case of SaFtsZ and HpFtsZ, but it did not have any effect on EcFtsZ. Further, the addition of PC190723 after the appearance of FtsZ polymers in yeast cells did not have any notable effect on the FtsZ structures **(Fig. S4)**. Our results presented here thus support the earlier reports that PC190723 acts to reduce the critical concentration of FtsZ polymerization of SaFtsZ (Andreu et al. 2010; Elsen et al. 2012).

Earlier studies had reported that PC190723 was non-toxic to eukaryotic cells, including budding yeast (Haydon et al. 2008). We further tested if PC190723 resulted in morphological defects in *S. pombe*, like sanguinarine and berberine, at higher concentrations. However, consistent with the earlier reports, PC190723 was inactive against *S. pombe* at both 56.2 μM and 140.6 μM and did not cause any morphological changes (**Fig. 2H iv**). Further, PC190723 did not disrupt the yeast microtubules at either of the concentrations **(Fig. S3 A iv and B iv)**.

### PC190723 stabilizes polymers and slows down the turnover rates of both SaFtsZ and HpFtsZ

We tested if PC190723 impacted the turnover rates of SaFtsZ and HpFtsZ using fluorescence recovery after photobleaching (FRAP) experiments. Fluorescence recovery was measured after bleaching a complete SaFtsZ spot or patch either in untreated cells (**Fig. 4A**) or in cells treated with 56.2 μM of PC190723 (**Fig. 4B**). Representative graphs showing the fluorescence recovery of SaFtsZ in the absence of PC190723 or the presence of 56.2 μM of PC190723 are depicted in **Figure 4C** and **Figure 4D**, respectively, and the plots for all such cells are shown **(Fig. S5 A and B)**. While SaFtsZ exhibited rapid dynamics with a half-time (*t_1/2_*) of 3.61 ± 0.41 seconds (95% CI; N = 10) in the absence of PC190723, the half-time (*t_1/2_*) of fluorescence recovery increased by nearly 3-fold to 11.34 ± 3.52 seconds (95% CI; N = 10) in the presence of PC190723 (**Fig. 4E**), giving an effect size of 7.7 [95% CI 4.0, 11.5]. Fluorescence recovery of HpFtsZ was similarly measured after bleaching a small part of HpFtsZ filaments in both untreated cells (**Fig. 4F**) or in cells treated with 56.2 μM of PC190723 (**Fig. 4G**). Representative graphs of the fluorescence recovery of HpFtsZ in the absence of PC190723 or the presence of 56.2 μM of PC190723 are shown in **Figure 4H** and **Figure 4I**, respectively. The plots for all such cells are shown **(Fig. S5 C and D).** The effect of PC190723 on the turnover rates of HpFtsZ was milder but significant and resulted in 1.96 times fold change in the fluorescence recovery half-time (*t_1/2_*) rates with an effect size of 5.9 [95% CI 3.5, 8.3]. The recovery half-time (*t_1/2_*) was 6.16 ± 0.89 seconds (95% CI; N = 10) in untreated cells and is consistent with the earlier reports (Specht et al. 2013). In the presence of PC190723, it increased to 12.06 ± 2.14 seconds (95% CI; N = 10) (**Fig. 4J**). Interestingly, compound 8j, a related benzamide derivative, has been shown to slow down FtsZ-ring turnover by 3-fold in *B. subtilis* (Adams et al. 2011). These results from the FRAP studies show that PC190723 affects the dynamics and turnover of both *S. aureus* and *H. pylori* FtsZ filaments, consistent with an effect of stabilization of filaments by the inhibitor.

**Figure 4.**
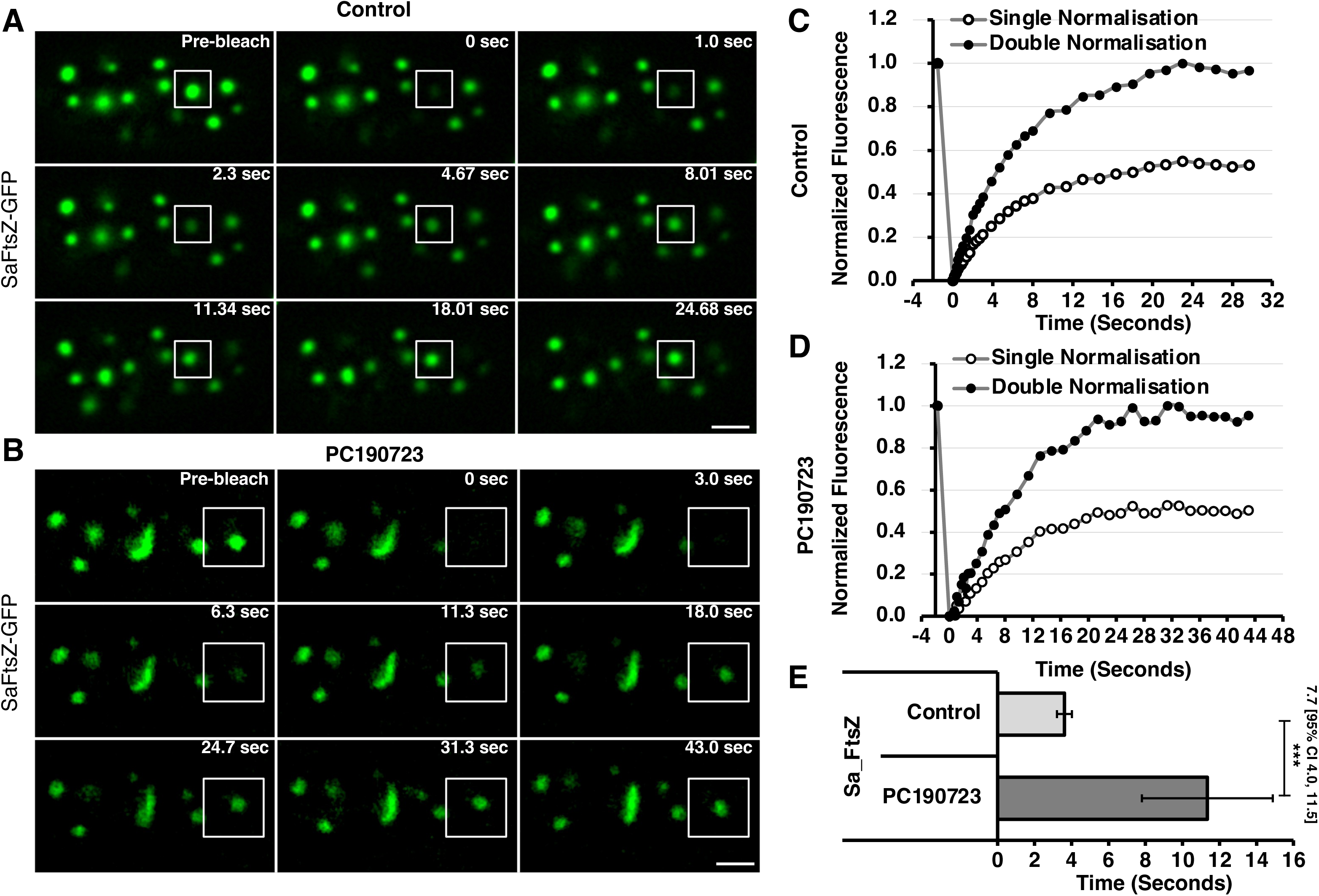
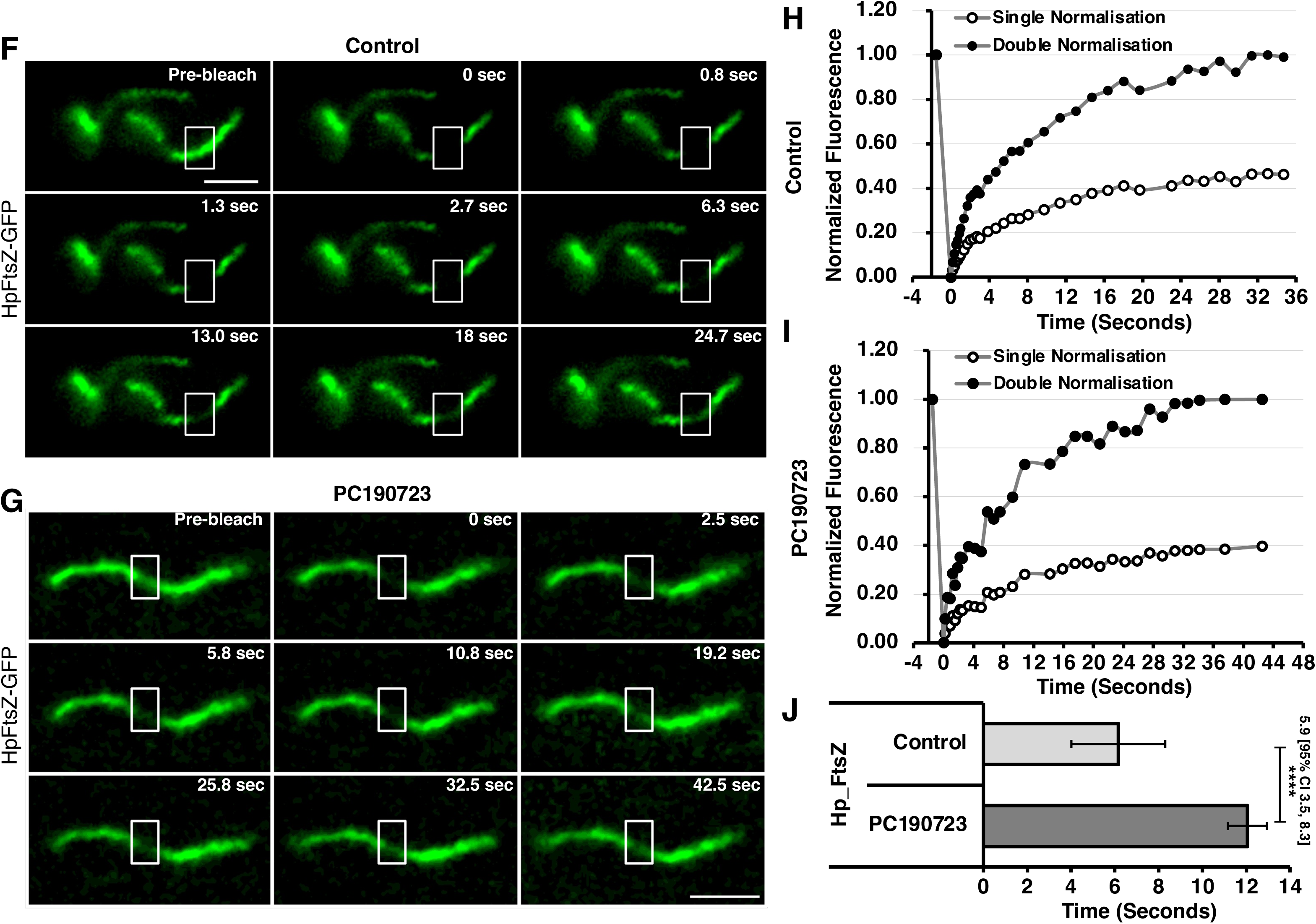
PC190723 reduces the polymer turnover rates of SaFtsZ and HpFtsZ. Fluorescence recovery after photobleaching (FRAP) of **(A)** control (DMSO treated) and **(B)** PC190723 (56.2 μM) treated cells expressing SaFtsZ. Fission yeast cultures expressing SaFtsZ-GFP were mounted onto agarose slides and imaged with an epifluorescence microscope. Fluorescence bleaching at selected ROIs were carried out using a 488 nm laser line. Fluorescence intensities during recovery was quantified, and a representative graph plotting the normalized intensities is shown in **(C)** for control (DMSO treated) cells and **(D)** PC190723 treated cells. **(E)** Quantification of the half-time (*t_1/2_*) values for control (N = 10) and PC190723 treated (N = 10) cells. Fluorescence recovery after photobleaching (FRAP) of **(F)** control (DMSO treated) and **(G)** PC190723 (56.2 μM) treated cells expressing HpFtsZ. Fission yeast cultures expressing HpFtsZ-GFP were imaged as mentioned above. Normalized fluorescence intensities of representative experiments are plotted for **(H)** control (DMSO treated) cells and **(I)** PC190723 treated cells. **(J)** Quantification of the half-time (*t_1/2_)* values for control (N = 10) and PC190723 treated (N = 10) cells. The *t_1/2_* values mentioned in the text were obtained by fitting the fluorescence recovery curves to a single exponential curve F(t)= C(1 − e^−kt^) using Excel’s Solver function and calculated from *t_1/2_* = ln(2)/k. An unpaired two-tailed Student’s t-test was used to assess the statistical significance. P-values ≤ 0.0001 and ≤ 0.001 are indicated by **** and ***, respectively. The mean difference between the proportion of cells carrying FtsZ polymers in the untreated and treated populations is also indicated as effect size [95% CI, lower bound, upper bound]. Scale bar represents 5 μm.

### SaFtsZ^G196A^ and HpFtsZ^G213A^ expressed in yeast exhibit PC190723 resistance

A comparison of the FtsZ sequences from various bacterial species that were tested for growth inhibition by PC190723 had revealed several key residues in FtsZ, determining its susceptibility or resistance (Haydon et al. 2008). In both *B. subtilis* and *S. aureus*, a single point mutation in FtsZ at G196 position to alanine (A) has been shown to result in a several-fold increase in the minimal inhibitory concentration of PC190723 (Stokes et al. 2013). Therefore, we next tested if we could experimentally observe resistance to PC190723 by FtsZ mutants G196A in our yeast expression system. Structure-based sequence alignment and comparisons showed that the equivalent residue of G196 in HpFtsZ is G213 and is conserved across the FtsZs **(Fig. S6)**. Hence we changed G213 in HpFtsZ to alanine (Ala) and expressed HpFtsZ^G213A^ in fission yeast. We then tested the effect of PC190723 on the mutation in a similar manner as described above, at a point when untreated cells predominantly exhibited diffuse fluorescence or short polymers.

In the case of HpFtsZ^G213A^, we found that only 41 ± 2.9 % (95% CI; N = 3; 500 cells were counted for each replicate) of the cells treated with 56.2 μM PC190723 as compared to 31 ± 16.4 % (95% CI; N = 3; 500 cells were counted for each replicate) in untreated cells had FtsZ polymers (**Fig. 5 A – C**). As compared to the effect size of 28.2 [95% CI 19.8, 36.7] for HpFtsZ^WT^, the effect size for HpFtsZ^G213A^ treated with PC190723 was 10.4 [95% CI −1.1, 21.9] with a negative value for the lower bound. Thus, there was no significant increase in the number of cells having FtsZ polymers in the mutant HpFtsZ^G213A^ upon treatment with PC190723 (**Fig. 5A**). In comparison, as described above, HpFtsZ^WT^ exhibited a significant increase in the number of cells having FtsZ polymers (70% of cells) when treated with PC190723 (**Fig. 5A and 5C**). Although PC190723 did not have an effect on the G213A mutant of HpFtsZ, HpFtsZ^G213A^ was proficient in polymerization and assembled into filaments **(Fig. S7A)**.

**Figure 5.**
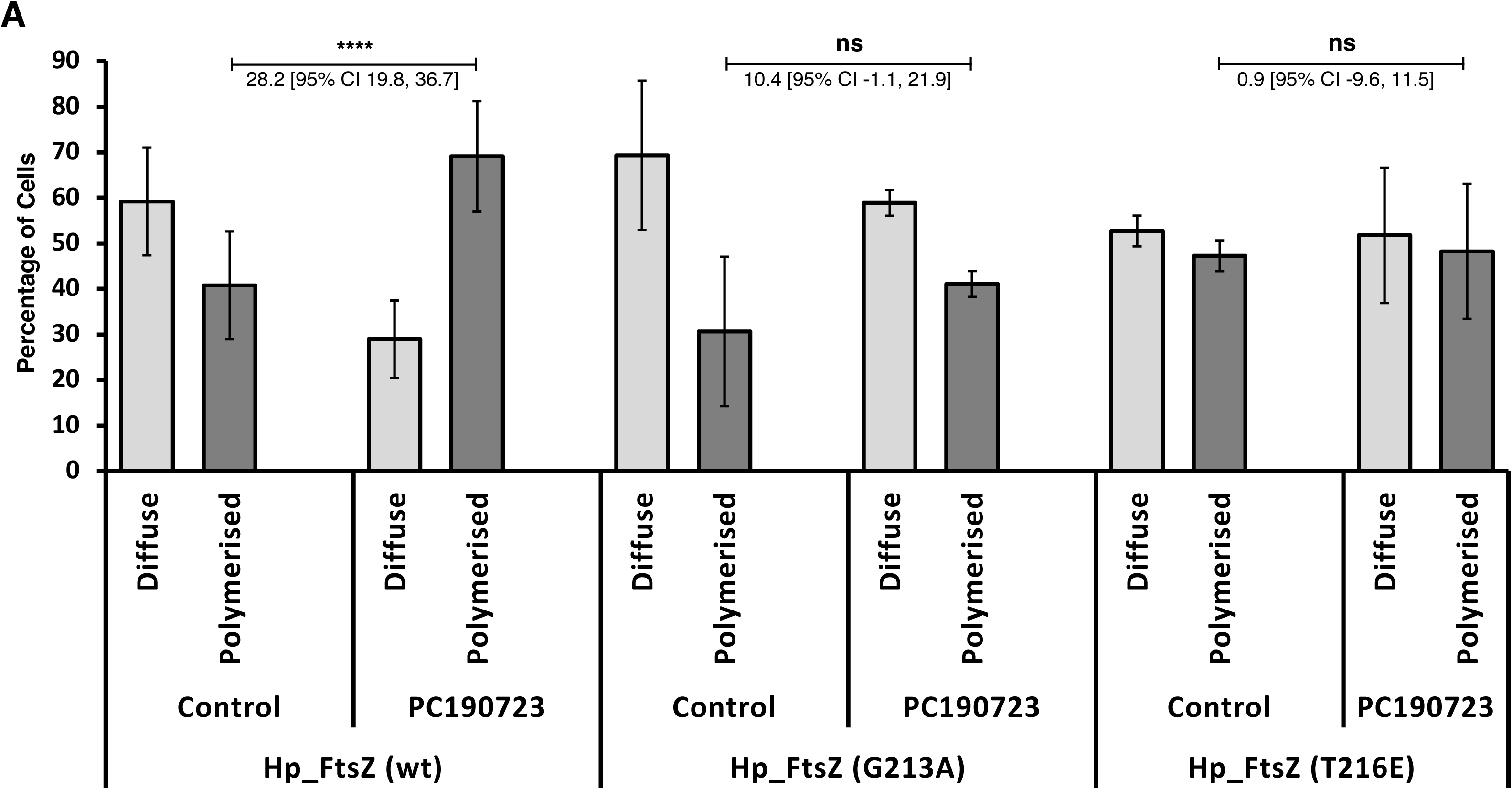
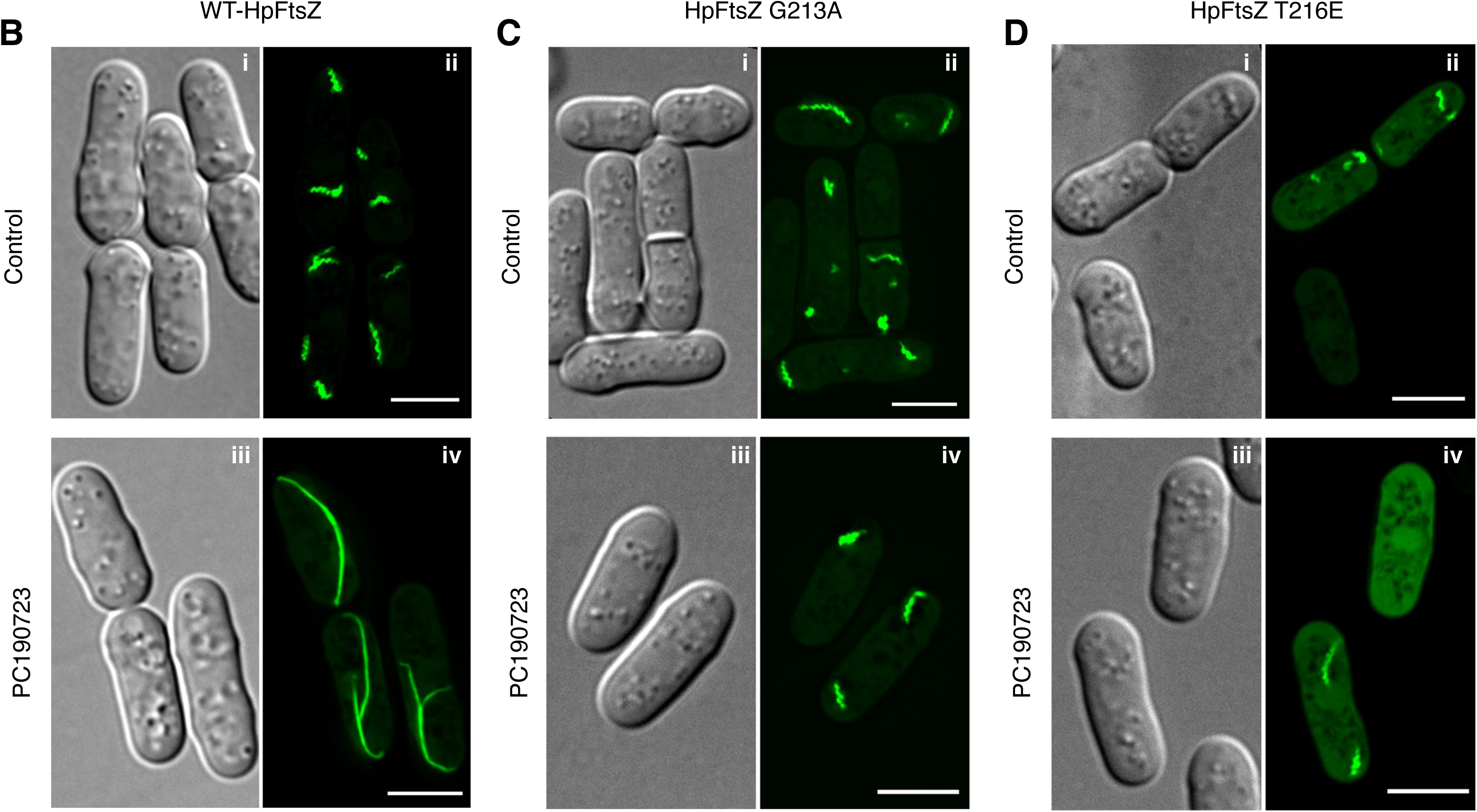
Determinants of resistance to PC190723 in HpFtsZ. *S. pombe* cultures carrying HpFtsZ^WT^, HpFtsZ^G213A^ or HpFtsZ^T216E^ were grown for 8 – 10 hours in the absence of PC190723 and then sub-cultured in the presence of the drug at 30°C for 4 hours to allow the sufficient expression of the proteins and imaged using an epifluorescence microscope. While G213A is predicted to act as a steric block to PC190723, T216E would result in salt-bridge interactions with R326 (R307 in EcFtsZ^WT^). **(A)** Effect of PC190723 on polymerization of HpFtsZ mutants G213A and T216E. The graph shows the percentage of cells having diffuse (light gray) and polymerized (dark gray) HpFtsZ in fission yeast in the absence (control) or presence of PC190723 (56.2 μM). Images showing the effect of PC190723 at 56.2 μM on **(B)** HpFtsZ^WT^ (panel iv), **(C)** HpFtsZ^G213A^ (panel iv) and **(D)** HpFtsZ^T216E^ (panel iv). At a time-point when control (DMSO treated) cells expressing HpFtsZ^WT^ show short polymers (panel ii), cells grown with PC190723 (56.2 μM) show stabilization of longer HpFtsZ polymers (panel iv). However, cells expressing **(C)** HpFtsZ^G213A^ (panel iv) and **(D)** HpFtsZ^T216E^ (panel iv) exhibit short polymers or diffuse fluorescence in the presence of 56.2 μM of PC190723 like in DMSO-treated cells expressing **(C)** HpFtsZ^G213A^ (panel ii) and **(D)** HpFtsZ^T216E^ (panel ii). The error bars shown are inferential and represent the 95% CI (N = 3). **** and ns indicate P-values ≤ 0.0001 and > 0.05 respectively. P-values were calculated from the two-sided proportions z-test. The mean difference between the proportion of cells carrying FtsZ polymers in the untreated and treated populations is also indicated as effect size [95% CI, lower bound, upper bound]. Scale bar represents 5 μm.

We also expressed SaFtsZ^G196A^ in fission yeast cells from the same *nmt41/42* promoter as for the wild-type SaFtsZ. Surprisingly, unlike HpFtsZ^G213A^, we found that SaFtsZ^G196A^ failed to assemble into any structures in yeast cells **(Fig. S7B ii)**. Although it has been shown that FtsZ^G196A^ is functional for cell division in both *S. aureus* and *B. subtilis* and forms Z-rings in the latter (Haydon et al. 2008; Stokes et al. 2013; Adams et al. 2016), mutations at G196 is also known to result in growth defects in *S. aureus* (Stokes et al. 2013; Adams et al. 2016). Nonetheless, the addition of 56.2 μM of PC190723 also did not induce any polymerization or the assembly of SaFtsZ^G196A^ **(Fig. S7B iv)**. Taken together, our results show that mutations that render FtsZ insensitive to PC190723 do not assemble early into polymeric structures in the presence of the drug, suggesting that PC190723 fails to bind SaFtsZ^G196A^ and HpFtsZ^G213A^.

### PC190723 docks into the inter-domain cleft of the HpFtsZ modelled structure in the T state

Crystal structures of SaFtsZ bound to PC190723 (**Fig. 6A and B**; PDB ID 4DXD) reveal the binding mode of PC190723 to the T state conformation of the protein (Matsui et al. 2012b; Tan et al. 2012). In the T state conformation of FtsZ, the cleft between the H7 helix and CTD is opened, and hence the inhibitor can be accommodated in the pocket. The residues Gly196, Leu200, Val203, Asn208, Leu209, Met226, Ile228, Asn263, Val297, Thr309 and Ile311 interact with the inhibitor in SaFtsZ (**Fig. 6A**). In the R state conformation of FtsZ, the inter-domain cleft is closed and hence does not have an adequate gap between the H7 helix and the central beta-sheet of CTD (S7, S8, S9, S10 strands) to accommodate PC190723 into the cleft. The inhibitor shows clashes with residues of the cleft when the T state structure (PDB ID: 4DXD) is superimposed with the R state structure (PDB ID: 5H5I) of SaFtsZ (**Fig. 6B**). A structure-based multiple sequence alignment reveals that many of the residues in FtsZ involved in the interaction with PC190723 are conserved in HpFtsZ **(Fig. S6 A and B)**.

**Figure 6.**
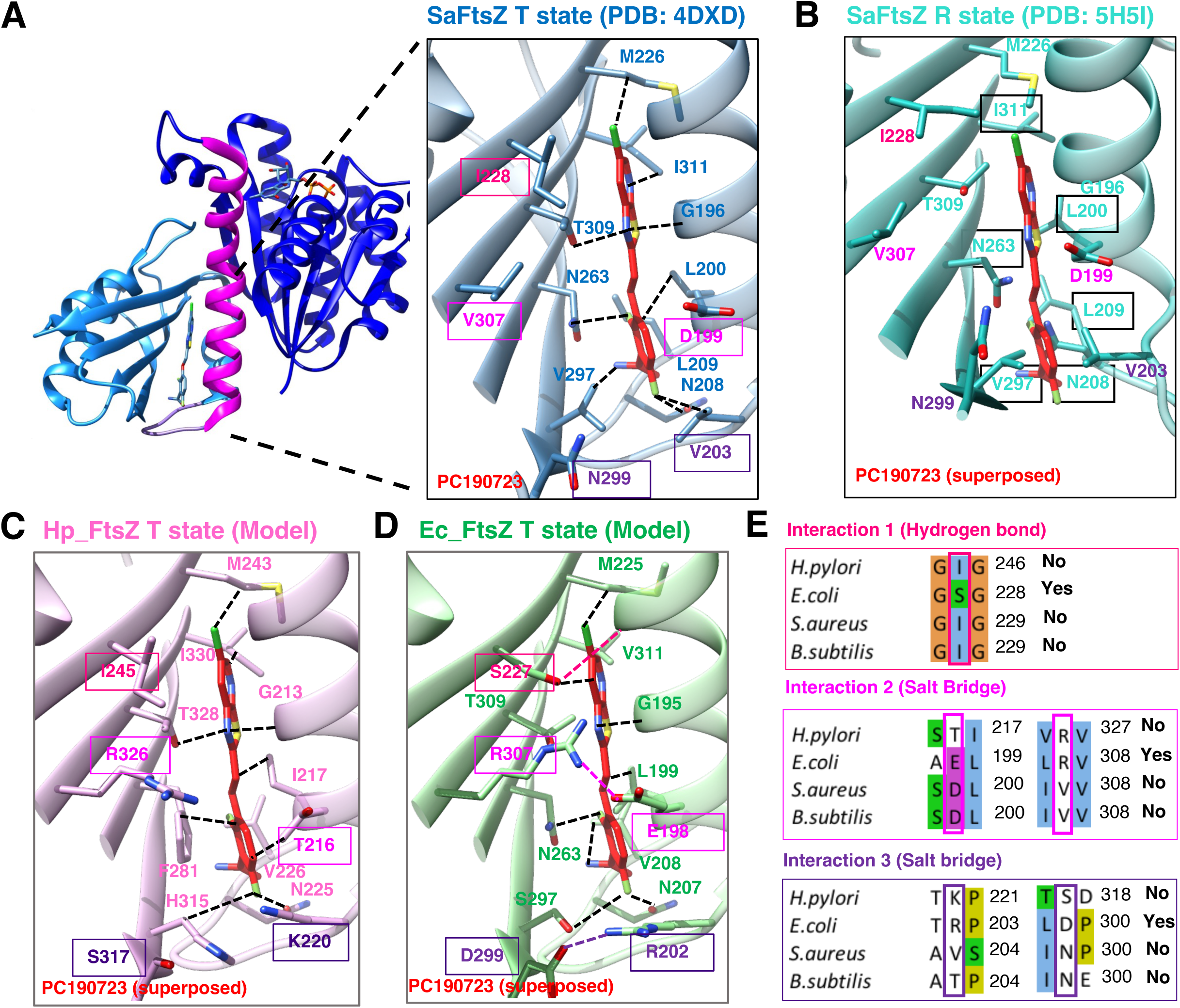
PC190723 can bind to the T state conformation of HpFtsZ. **(A)** Overall structure of SaFtsZ bound to PC190723 (PDB ID: 4DXD; NBD - dark blue, H7 helix-magenta, T7 loop-purple and CTD-blue) with a zoomed-in view of PC190723 binding cleft (PDB ID: 4DXD; T state, open cleft) as inset. Residues of FtsZ, which interact with PC190723 (PDB ID-4DXD), are shown in stick representation. **(B)** PC190723 binding cleft in R state conformation (PDB ID: 5H5I; sea green) is closed, and the residues clash with the drug (highlighted in black boxes) and prevent the binding of the drug. **(C)** HpFtsZ modelled in T state structure (light pink) and superimposed on PDB 4DXD. There are no clashes with the drug, and F281 could stack against the benzamide ring of PC190723. HpFtsZ does not possess any interaction that could potentially block the cleft. **(D)** PC190723 placed (by superposition of PDB 4DXD onto EcFtsZ T state model) in the pocket in EcFtsZ modelled in T state structure (green) highlight that there are two salt bridges, R307 and E198; D299 and R202 (shown in magenta and purple respectively) and a hydrogen bond between OH group of S227 and the main chain of G191 in H7 helix (shown in dark pink) which could block the cleft. **(E)** Residues in HpFtsZ and SaFtsZ corresponding to residues of EcFtsZ, which might block the PC binding cleft due to the three interaction types highlighted in the figure panels. In all panels, residues which are labelled in dark pink, magenta and purple fonts and boxed are the corresponding residues which form a salt bridge and hydrogen bonds in EcFtsZ. Dotted lines represent interactions at distances ≤ 4Å.

We further resorted to *in silico* approaches to determine if the inter-domain cleft in HpFtsZ could accommodate PC190723. We thus modelled HpFtsZ using SwissModeller (Waterhouse et al. 2018) with the T state of SaFtsZ as a template and superposed it with the PC190723 bound SaFtsZ structure (4DXD) to see whether PC190723 can fit into the modelled HpFtsZ structure (**Fig. 6C**). Analysis of PC190723 in the inter-domain cleft of the HpFtsZ model indicated that several residues that made direct contact with the drug in the crystal structure of the PC190723-SaFtsZ complex (4DXD) were conserved in HpFtsZ and shown to be within interacting distances of PC190723. Phe281 of HpFtsZ could potentially interact by a stacking interaction with the ring of PC190723. The side chain of Thr328 is at a distance capable of forming hydrogen bonds, while the side chain of Ile217 forms van der Waals interaction with the inhibitor (**Fig. 6C and Fig. S6B)**. Unlike in the R state, the residues in the S7, S8, S9 and S10 beta-sheets or the H7 helix do not clash with the inhibitor in the T state.

### Resistance to PC190723 is mediated by salt-bridge interactions that prevent drug binding in the inter-domain cleft

A single point mutation in FtsZ at position Gly196 in *B. subtilis* and *S. aureus* is sufficient to render them resistant to PC190723. However, it was also noted that while the susceptible species were characterized by the presence of valine (Val) at position 307, some other species of bacteria that were naturally resistant to PC190723 contained arginine (Arg) or histidine (His) at this position (Haydon et al. 2008; Kaul et al. 2013b). Surprisingly, HpFtsZ contains an arginine at this position (Arg326), and yet our results from the yeast expression studies suggested that PC190723 had an effect on the polymerization and dynamics of HpFtsZ. In order to further probe this apparent discrepancy, we analyzed the sequences and molecular structures of FtsZ from various species that included both susceptible and resistant species **(Fig. S6A)**. A detailed structure-based sequence analysis of FtsZ showed that in species that are resistant to PC190723, there are possibly two salt bridges that might play an important role in occluding the inter-domain cleft (**Fig. 6D and Fig. S6)**. The blockage of this cleft might prevent the binding of the inhibitor (**Fig. 6D**). In species like *E. coli* that are resistant to PC190723, these blockages constitute two salt bridge interactions-Arg307 and Glu198 (shown in red); Asp299 and Arg202 (shown in violet) and one hydrogen bond interaction between the OH group of Ser227 and the main chain of Gly191 (shown in brown) (**Fig. 6D**). These interactions are absent in *S. aureus* FtsZ; thereby, the inhibitor can be accommodated in the cleft. However, in HpFtsZ, the equivalent position of Glu198 in EcFtsZ is threonine (Thr216), which is smaller than glutamic acid (Glu) and probably far off to engage in any interaction with Arg326 (the equivalent of Arg307 in EcFtsZ) (**Fig. 6E**). Thus, the salt bridge which might block the inhibitor from binding at the cleft is not present in HpFtsZ (**Fig. 6E**), thereby possibly allowing PC190723 to bind.

Therefore, restoring the salt-bridge interactions between the amino acids at positions 326 and 216 in HpFtsZ should render it insensitive to PC190723. In order to test this hypothesis, we changed Thr216 in HpFtsZ to glutamic acid (Glu), expressed the mutant HpFtsZ^T216E^ in fission yeast and tested the effect of PC190723 on this mutation. In accordance with the structural analysis, we found no significant difference in the number of cells having FtsZ polymers in the mutant HpFtsZ^T216E^ upon treatment with PC190723 (**Fig. 5 A and D**). In the presence or absence of PC190723, HpFtsZ^T216E^ expressing cells showed 47 ± 3.4 % (95% CI; N = 3; 500 cells were counted for each replicate) and 48 ± 14.8 % (95% CI; N = 3; 500 cells were counted for each replicate) respectively, that assembled FtsZ polymers (**Fig. 5A**). This was unlike HpFtsZ^WT^ (**Fig. 5A and 3D**), where almost 70 % of the cells treated with PC190723 contained polymers as compared to a mere 38 % in untreated cells (**Fig. 5A and 3D**). The effect size for HpFtsZ^WT^ treated with PC190723 was 28.2 [95% CI 19.8, 36.7] as compared to the effect size for HpFtsZ^T216E^ treated with PC190723, which was 0.9 [95% CI −9.6, 11.5] with a negative value for the lower bound of the confidence interval. Further, like HpFtsZ^G213A^, HpFtsZ^T216E^ was capable of polymerization and assembled into polymers upon expression in fission yeast **(Fig. S7C)**. Our results here, therefore, suggest that the salt-bridge interactions between residues in the H7 helix and adjacent sheets (S9 and S10) in the inter-domain cleft form the basis for the inability of FtsZ in many species to bind to PC190723 rather than a single residue (Arg or His) at position 307 (EcFtsZ).

## Discussion

In this study, we have attempted to develop a cell-based assay using fission yeast (*S. pombe*) as a heterologous expression host, which would enable the screening of compounds that could directly affect FtsZ polymerization as well as identify potential toxicity to yeast (or eukaryotic) cells simultaneously. As a proof-of-principle, we first assessed the ability of FtsZ from two pathogenic species, *S. aureus* and *H. pylori,* to assemble into structures in fission yeast and tested the effects of known FtsZ modulators, sanguinarine, berberine and PC190723 on these FtsZ assemblies. We show that both SaFtsZ and HpFtsZ assembled into structures in fission yeast. While SaFtsZ assembled into spots/ patches or ring-like structures, HpFtsZ assembled into both linear and spiral polymers. Interestingly, spiral filaments of FtsZ polymers of *H. pylori* have also been reported in bacterial cells and when expressed in *Drosophila melanogaster* S2 Schneider cells (Specht et al. 2013), suggesting that assembly into helical filaments might be intrinsic to HpFtsZ polymers. Previous studies have shown that various factors such as molecular crowding, variable C-terminal regions and bound nucleotide state lead to the formation of supramolecular structures like twisted helical structures, toroids and rings similar to those that have been observed *in vivo* (Popp et al. 2009; Huecas et al. 2017). Thus, the molecular crowding due to the dense cytoplasm of the yeast cells could have possibly induced the spiral and ring-like assembly of FtsZ polymers (Erickson et al. 2010).

Several approaches have been used to screen small molecules targeting bacterial cell division and FtsZ. While *in vitro* methods such as NMR (Domadia et al. 2007; Sun et al. 2014; Araújo-Bazán et al. 2019) and crystallography (Läppchen et al. 2008; Fujita et al. 2017) are valuable and offer information on distinct binding sites, these are not efficient for screening. Electron microscopic examination can distinguish the effects of the compounds being tested on the FtsZ protofilament assembly and lateral associations (Nova et al. 2007; Kaul et al. 2012; Anderson et al. 2012; Sun et al. 2014; Huecas et al. 2017; Kumar et al. 2011; Park et al. 2014). Other techniques that are routinely used include fluorescence anisotropy (Ruiz-Avila et al. 2013; Park et al. 2014), 90° light-scattering assay (Mukherjee and Lutkenhaus 1999) and dynamic light scattering (Hou et al. 2012; Di Somma et al. 2020) for assessing inhibition of FtsZ assembly (Irwin et al. 2015; Lui et al. 2019; Kaul et al. 2012; Nova et al. 2007; Anderson et al. 2012). Other easily scalable high-throughput assays include FCS/FCCS and FRET-based methods (Hernández-Rocamora et al. 2015; Mikuni et al. 2015; Reija et al. 2011).

*In vivo* assays relying on cell filamentation phenotype coupled with the localization of Z-ring might be a good indicator of FtsZ being the direct target. However, since bacteria can undergo cell filamentation and not assemble FtsZ rings in response to a variety of conditions, including DNA damage (Mukherjee et al. 1998) and disruption of membrane potential (Strahl and Hamoen 2010), the *in vivo* assay is not so useful unless combined with the *in vitro* assays mentioned above. Finally, the isolation of resistance mutants in FtsZ to the drug can provide strong evidence of FtsZ being the direct target.

Reconstitution systems are powerful and provide excellent control over the system, but they are emerging technologies and are technically challenging. Reconstitution systems include a variety of methods, such as the use of membrane nanodiscs, microbeads of different materials, supported bi-layer membranes (SLBs) and biomimetic systems that provide cell-like environments (Monterroso et al. 2013; Rivas et al. 2014). While *in vitro* biochemical assays and reconstitution systems are useful to find molecules that directly target FtsZ, they are cumbersome and need to be performed at optimal physiological pH and ionic conditions, which can be considerably variable among FtsZ from different species.

Our results on the ability of sanguinarine and berberine to specifically affect the assembly of FtsZ and not MreB in fission yeast highlight the utility of the heterologous expression system as a platform to identify molecules that specifically affect FtsZ polymerization. The yeast platform offers a cellular context mimicking the cytoplasm for cytoskeletal assembly. The system is simple to replicate in any laboratory, including those focused on chemical synthesis with minimum microbiological expertise and can be easily reproduced and scaled up as well. However, one of the major disadvantages of using fission yeast could be the need to use much higher concentrations of drugs than normally used for mammalian cell cultures to achieve an inhibitory effect. This could probably be due to the poor permeability of certain drugs in fission yeast because of its thick cell wall (Benko et al. 2017; Pérez and Ribas 2004). A similar effect of toxicity might arise at much lower concentrations in other eukaryotic cells, such as human cells. Consistently, while sanguinarine and berberine are known to affect the eukaryotic microtubules at 10 μΜ – 20 μM concentrations (Lopus and Panda 2006; Wang et al. 2016; Raghav et al. 2017), morphological effects on yeast cells were observed only at concentrations > 100 μM. However, yeast microtubules were not affected by berberine and sanguinarine. Differences in membrane lipid profiles and MDR efflux pumps between yeasts and mammalian cells might also contribute to differential resistance to the drugs being tested (Balzi and Goffeau 1991). Conversely, an inhibitory effect in yeast cells may not necessarily translate into toxicity in a human cell. These and the permeability of drugs in yeast cells represent an important caveat in using such heterologous expression systems for the screening of compounds against target molecules. However, notwithstanding this caveat, the heterologous system provides significant advantages in assessing the direct effects of the drug on FtsZ assembly. Moreover, fission yeast-based high-throughput platform screening methods using imaging have been successfully adapted to the screening of drugs against HIV-1 proteases by large-scale screening facilities such as the NIH Molecular Libraries Probe Production Centers Network in the Molecular Libraries Program, leading to several candidate drugs (Benko et al. 2017, 2019). This aspect also emphasizes the benefit of utilizing an economic eukaryotic heterologous host for testing drugs targeting bacterial proteins. Any compound that potentially affects eukaryotic function can be easily identified and discarded at the screening step, thus reducing time and costs on further pharmacological analysis in the drug development pipeline.

PC190723 has been one of the most promising drugs against FtsZ and remains the only inhibitor for which the binding site in FtsZ has been clearly established. However, PC190723 predominantly affects Gram-positive bacteria and FtsZ from Gram-negative bacteria have been shown or predicted to be resistant (Haydon et al. 2008). Several efforts have been made to target Gram-negative bacteria with derivatives of benzamide. Examples include difluorobenzamides, substituted benzodioxanes, heterocyclic and non-heterocyclic derivatives (Straniero et al. 2017; Chai et al. 2020; Straniero et al. 2020a, 2020b). Although many exhibited promising activity *in vitro*, most were substrates for the AcrAB class of efflux pumps (Chai et al. 2020; Kaul et al. 2014; Straniero et al. 2020a, 2020b; Casiraghi et al. 2020). Thus, the poor membrane permeability, signature outer membrane, particularly lipopolysaccharide (LPS) structure (Wang et al. 2021), the presence of multiple efflux pumps in species such as *E. coli*, *Klebsiella pneumonia* and *Pseudomonas aeruginosa* (Piddock 2006), and differences in FtsZ sequences in the binding-site (Kaul et al. 2013b; Miguel et al. 2015) have been cited as reasons for lack of susceptibility of Gram-negative bacteria to benzamide derivatives (Casiraghi et al. 2020). More recently, two molecules, TXA6101 and TXY6129, with substituted 2,6-difluorobenzamide scaffold, have been shown to inhibit the polymerization of both *E. coli* and *Klebsiella pneumoniae* FtsZ. Moreover, despite being substrates for efflux pumps, TXA6101 induced morphological changes in K. pneumoniae (Rosado-Lugo et al. 2022). Studies in the past on the effects of PC190723 on *E. coli* have been confusing, with a few reports suggesting an effect on FtsZ polymerization resulting in cell filamentation (Kaul et al. 2014), while others did not find any effect on EcFtsZ (Andreu et al. 2010; Anderson et al. 2012; Khare et al. 2019). The outer membrane has been shown to be a permeability barrier for PC190723 in *E. coli* (Khare et al. 2019; Chai et al. 2020). In addition, the Resistance-Nodulation-Division (RND) family of efflux pumps has been attributed to resistance against 2,6-difluorobenzamide derivatives, including TX436 (a prodrug of PC190723) in Gram-negative bacteria (Kaul et al. 2014).

It was, therefore, of interest to test the effects of PC190723 on FtsZ assembly in the yeast expression system, wherein the interpretations are not affected by the compounding factors such as differential membrane permeability or efflux pumps. Our assay provides for a direct comparison of the effect of PC190723 on FtsZ from different species. While our results presented here clearly suggest that PC190723 does not affect the assembly of EcFtsZ, it exerted an effect on HpFtsZ, which is also from a Gram-negative bacterium. Assemblies of SaFtsZ and HpFtsZ structures appeared in fission yeast cells at a much earlier time when treated with PC190723, suggesting that the drug acted to lower the critical concentration of FtsZ polymerization. Further, FRAP experiments performed on SaFtsZ and HpFtsZ suggest that PC190723 affects the dynamics of FtsZ polymers and slows down the turnover rates. We had earlier shown that FtsZ mutants defective in GTP hydrolysis expressed in fission yeast have slower turnover rates, as measured by fluorescence recovery half-times (*t_1/2_*) rates (Srinivasan et al. 2008). These results are also consistent with the earlier findings that PC190723 acts to induce FtsZ polymerization and stabilize FtsZ filaments (Andreu et al. 2010; Elsen et al. 2012; Miguel et al. 2015; Fujita et al. 2017) and its derivative compound, 8j acting to slow down FtsZ-ring turnover by 3-fold in *B. subtilis* (Adams et al. 2011).

Finally, our studies here suggest that resistance to PC190723 in FtsZ from Gram-negative bacteria like *E. coli* is a result of occluding salt-bridge interactions in the inter-domain cleft between the H7 helix and adjacent sheets (S9 and S10) and may not be due to a single residue (Arg or His) at position 307. Although HpFtsZ contains an arginine at the equivalent position (Arg326), this residue cannot engage in a salt bridge since HpFtsZ has a threonine (Thr) at position 216 (the equivalent of Glu198 in EcFtsZ). Restoring the salt-bridge interaction in HpFtsZ by changing Thr216 to glutamate (E) conferred resistance to PC190723 in our yeast expression studies. However, PC190723 failed to exert growth inhibition of *H. pylori,* probably owing to outer membrane permeability barriers. Thus, although further genetic, cell biological and biochemical studies will be required to fully elucidate the resistance of *H. pylori* cells to PC190723, our studies here provide new insights into the molecular determinants in FtsZ that lead to resistance against PC190723. A powerful emerging technique based on cytological profiling has been successfully used to identify the cellular pathways targeted by the inhibitors (Nonejuie et al. 2013; Martin et al. 2020), including cell division inhibition by FtsZ (Araújo-Bazán et al. 2016). The recent advances in computational image analysis and deep learning approaches (von Chamier et al. 2021; Spahn et al. 2022) could further advance image-based screening for FtsZ inhibitors (Andreu et al. 2022). Thus, our results here suggest that the fission yeast expression system can be used to identify small molecules that directly target FtsZ, probe the molecular determinants of resistance and establish a proof-of-concept of an *in vivo* cell-based assay to discover antimicrobials that specifically target the bacterial cytoskeleton.

## MATERIALS AND METHODS

### Growth and culture of bacteria and yeast

The strains and plasmids used in this study are listed in **Supplementary Table S1**. *E. coli* strains and *S. aureus* (ATCC 25923) were grown in LB medium at 37°C, whereas *H. pylori* (strain *H. pylori* 26695) was grown in Brucella broth with 10 % fetal bovine serum media at 37°C with 10 % CO_2_. *E. coli* cultures carrying plasmids were grown in LB medium containing appropriate antibiotics (carbenicillin at 100 μg/ mL; kanamycin at 50 μg/ mL; chloramphenicol at 34 μg/ mL). *S. pombe* strain 192 (*h*− *leu1-32 ura4-D18*) and mCherry-Atb2 strains were grown at 30°C in YES medium. Transformation of *S. pombe* was carried out using the Lithium acetate method as described previously (Bähler et al. 1998) and plated on selective Edinburg minimal medium (EMM) plates containing 15 μM thiamine. EMM plates and broth were supplemented with adenine (0.225 mg/ mL of media), histidine (0.225 mg/ mL) and leucine (0.225 mg/ mL) or uracil (0.1125 mg/ mL) as necessary. Thiamine was used at a final concentration of 15 μM on the agar plates to repress expression from the medium-strength pREP41/42 plasmids. All experiments were done with *S. pombe* growing at 30°C in exponential phase (OD_600_ of 0.3 – 0.6) and were maintained thus by intermittent dilution of cultures. Inhibitors PC190723, sanguinarine, berberine and A22 were dissolved in DMSO and added to *S. pombe* cultures at final concentrations as indicated in the text or figure legends.

### Plasmids and constructs

#### Cloning of EcFtsZ, HpFtsZ & SaFtsZ into yeast expression vector pREP42

EcFtsZ, SaFtsZ and HpFtsZ refer to FtsZ from *E. coli*, *S. aureus* and *H. pylori*, respectively. The cloning of EcFtsZ-GFP (pCCD4) into pREP42 has been described previously (Srinivasan et al. 2008). Genomic DNA was isolated from *H. pylori* and *S. aureus* wild-type strains using the Genomic DNA Isolation kit. These genomic DNA were used as a template in a PCR reaction using high-fidelity Q5 polymerase to amplify the *ftsZ* genes of these bacteria. The plasmids pCCD712 and pCCD713 expressing *H. pylori* FtsZ-GFP (HpFtsZ-GFP) and *S. aureus* FtsZ-GFP (SaFtsZ-GFP), respectively, were constructed by cloning the amplified PCR product into pCCD51 using restriction-free (RF) cloning method (van den Ent and Löwe 2006). pCCD51 was constructed as follows: GFP was amplified by PCR from pCCD4 using oligonucleotides RSO73 and RSO74. The PCR product was subsequently digested with SmaI and ligated into pREP42 (Moreno et al. 2000), linearized with the same enzyme. Site-directed mutagenesis was carried out following the Stratagene™ Quickchange-II site-directed mutagenesis procedure using high-fidelity Q5 polymerase. The oligonucleotides used are listed in **Supplementary Table S2,** and the list of chemical reagents with their catalogue numbers is listed in **Supplementary Table S3**.

### Microscopy and Fluorescence Recovery after Photobleaching

For imaging, the cells were mounted onto 1.7% or 2% agarose pads containing media in cavity slides and covered with a coverslip No. 1.5 prior to sealing with VALAP (1:1:1 mixture of vaseline, lanolin and paraffin). Images were acquired using an epifluorescence microscope (DeltaVision™ Elite) with a 100 X oil immersion objective lens of NA 1.4 and equipped with a Photometrics CoolSNAP™ HQ2 camera. An oil with a refractive index of 1.516 was used for imaging yeasts. Excitation filters and emission filters of 475/28 nm and 525/48 nm were used to image GFP. Z-stacks were acquired with a Z-depth of 0.2 or 0.3 μm. SoftWorX™ software was used to perform deconvolution using the iterative-constrained algorithm (Agard 1984) with the number of cycles set to 10 (ten). 3D-SIM on *S. pombe* was performed as described previously (Pande et al. 2022). Cells were mounted on an agarose pad as described above and imaged using a DeltaVision OMX-SR Blaze microscope. Raw images were acquired using a 60×, NA 1.42 oil-immersion objective lens and a PCO Edge 4.2 sCMOS camera. An exposure time of 2 – 5 ms was used to acquire each image, and Z-stacks were obtained at a step size of 0.125 μm to cover the width of *S. pombe* cells. 3D-SIM were reconstructed using the SI reconstruction module of SoftWorx™ software. FRAP experiments were carried out essentially as described before (Recovery half-times were calculated as described previously (Anderson et al. 2004; Srinivasan et al. 2008) except that a DeltaVision™ equipped with a Quantitative laser module (QLM; X4 Laser) was used for imaging. Photobleaching was carried out using 80% laser power with an argon laser (488 nm) with a pulse time of 0.05 sec. Images were acquired before and after photobleaching, and Z-stack images were acquired at fixed intervals with a Z-depth of 0.2 or 0.3 μm. Time-lapse intervals were varied or were typically 1 – 2 seconds apart and acquired for over 2 minutes. Integrated fluorescence intensities in the pre-bleached images and at each time point in the bleached and other regions were measured in Fiji (v 2.0.0-rc-69/1.52p). FRAP experiments were performed on different days, and data were pooled for analysis. All values were exported to Excel (Microsoft™) for further analysis. The recovery half-time (*t_1/2_*) values mentioned in the text were obtained by fitting the fluorescence recovery curves to a single exponential curve F(t)= C(1 − e^−kt^) and calculated from *t_1/2_* = ln(2)/k. The solver function in Excel was used for the least-squares fit of the corrected fluorescence intensities over time to the single exponential equation. All images were processed using Fiji (v 2.0.0-rc-69/1.52p) (Schindelin et al. 2012).

### Quantitation of the number of spots, polymer length and density

Cells were processed as described above for imaging, and images were acquired as described previously (Pande et al. 2022) for the calculation of the number of spots, polymer length and density. Briefly, imaging was carried out with constant parameters such as fixed illumination of 2% transmission (ND filter), exposure of 0.25 s, binning (1 x 1) and camera gain of 0.5 x). Both differential interference contrast (DIC) and fluorescence channels were imaged, and cell outlines were marked using the freehand drawing tool in Fiji (Schindelin et al. 2012). The cell outlines were saved as regions of interest using the ROI manager. Masking of areas outside of the cell outlines was carried out as described (Higaki 2017) and filled with black colour. The Number of spots per cell was then measured using the OPS threshold IJ1 analyze macro, which is available as a built-in feature in Fiji (v2.0.0-rc-69/1.52p; Schindelin et al. 2012). The macro utilizes a convolve filter with a 5 x 5 kernel and the ‘otsu’ method for auto-thresholding. A convex hull is used to create the masks of segmented particles, following which the ‘analyze particles’ plugin is run by the macro. For measuring polymer length and density (amount of cytoskeleton per unit area in a cell), images were processed with the inclusion of masking and skeletonization steps as previously described (Higaki 2017; Henty-Ridilla et al. 2014). Skeletonization was carried out by applying the lpx 2Dfilter in the lpx plugin (Higaki 2017). The specific parameter values giwsIter, mdnmsLen, pickup, aboveThr, shaveLen, delLen, draw, and preGauss were fixed as 5, 15, otsu, default, 5, 5, orgInt and −1, respectively, in the linemode under LineExtract. The LineFeature outputs the measure of the number of pixels with intensities (nPix) after skeletonization and the total polymer length within the selected ROIs. Density, the amount of cytoskeleton assembled per unit area in a cell, is then calculated as nPix divided by the area of the cell. The area of the cell is obtained using the measurement tool in the ROI manager from Fiji.

### Sequence and structural analysis for PC190723 binding to FtsZ

The sequences of FtsZs from various organisms were taken from UniProt, aligned using Clustal Omega (Sievers et al. 2011), and represented using Jalview (Waterhouse et al. 2009). The reference structure taken for the analysis was the previously reported T state structure of SaFtsZ bound to PC190723 (PDB ID: 4DXD), and the residues around 4 Å were determined using UCSF Chimera version 1.13.1 (Pettersen et al. 2004). The residues in SaFtsZ, which contributed to the binding of the inhibitor, were highlighted in the sequence alignment. The reference R state structure (PDB ID: 5H5I) was chosen to see whether the inhibitor can be accommodated in the R state structure of SaFtsZ by superimposing the C-alpha atoms of the nucleotide-binding domain (NBD) of PDB IDs: 5H5I and 4DXD. FtsZs from *H. pylori* and *E.coli* were modelled in T state using SWISS-MODEL (Waterhouse et al. 2018), with PDB ID 4DXD as the template. Similarly, the NBD of the modelled structures was superimposed onto the SaFtsZ structure bound to PC190723 and checked whether the inhibitor could be accommodated in the corresponding pocket of the models without significant clashes.

## Statistical Analysis

All statistical analysis and tests were carried out in Excel (Office 365 or Excel for Mac 2011) using the built-in Data Analysis functions or the add-ins XLSTAT or Xrealstat (Real Statistics). A single colony of bacteria or a patch of yeast cells independently grown from freshly streaked out cells on agar plates from frozen glycerol stocks was considered to be a biological replicate (N). A minimum of at least 3 biological replicates was used for every experiment. The dosage effect upon increasing concentrations of PC190723 on FtsZ assembly was assessed using the Cochran-Armitage test. A two-proportions z-test was used for the statistical analysis of the effect of PC190723 on the proportions of cells carrying FtsZ polymers and the calculation of P-values. An unpaired two-sided Student’s t-test was used as a statistical test for the FRAP experiments. Statistical tests and P-values are indicated in the figure legends using an asterisk. A P-value ≤ 0.0001, ≤ 0.001, ≤ 0.01, < 0.05 or ≥ 0.05 indicated by ****, ***, **, * and ns respectively. As an alternative to null-hypothesis significance tests (NHST), we also used estimation statistics to determine the effect size and 95 % confidence intervals (Claridge-Chang and Assam 2016; Ho et al. 2019). Superplots were plotted as described (Lord et al. 2020) using Excel Office 365. The error bars plotted are inferential and represent the 95% CI (Cumming et al. 2007; Vaux 2014; Pollard et al. 2019).

## Acknowledgements

The authors are grateful to Prof. Jose Manuel Andreu and Dr. Lidia Araujo-Bazán for their critical reading of the article and valuable comments and suggestions. The authors thank Dr. Asima Bhattacharyya (SBS, NISER, India) for *H. pylori* genomic DNA and Dr. Mithilesh Mishra (DBS, TIFR, Mumbai, India) for strains and plasmids. This work is supported by the Department of Biotechnology (DBT) from BT/PR15183/BRB/10/1443/2015 to RS and BT/PR42977/BRB/10/2022/2021 to RS & GP and DBT Membrane Structural Biology Program grant - BT/PR28833/BRB/10/1705/2018 to PG. PG also acknowledges support from INSPIRE Faculty Fellowship from (IFA12/LSBM-52), Department of Science and Technology, Innovative Young Biotechnologist Award (BT/07/IYBA/2013), Department of Biotechnology (DBT) Membrane Structural Biology Program grant (BT/PR28833/BRB/10/1705/2018), and Science and Engineering Research Board (SERB) Core Research grant (CRG/2018/003795) for infrastructure and staff support. Intramural core funding from the Department of Atomic Energy (DAE) and support staff members at the Centre of Interdisciplinary Sciences, NISER, are acknowledged. We also acknowledge fellowships from DAE to AKS, SMP, BSN, SK, NM and from IISER Pune to JC.

## Author contributions

AKS, SMP, BSN, JC and NM conducted the experiments. RS, GP and JC designed and analyzed the work. BSN initiated the work, SMP contributed to initial optimization, and AKS optimized further and carried out the experiments. AKS carried out the all analysis and quantification of the data with respect to the effect of inhibitors on FtsZ assembly. SMP carried out work related to 3D-SIM imaging, quantification, the resistance mutants and the effect of inhibitors on yeast microtubules. JC carried out all the *in silico* work and structural analysis. AKS, JC, GP and RS wrote the manuscript. AKS, SMP and JC edited the draft manuscripts. SK contributed to the plotting and analysis of FRAP data. NM carried out experiments related to MreB and A22 treatment.

## Notes

### Competing Interest Statement

The authors have declared no competing interest.

### Summary of Updates

The whole manuscript has been revised to address the reviewers comments and Figures 1 and 2 of the earlier version have now been deleted. New quantitative data has been added to figures 2 and 3 of the current version.

## References

Adams DW, Wu LJ, Czaplewski LG, Errington J. 2011. Multiple effects of benzamide antibiotics on FtsZ function. Mol Microbiol 80: 68–84.

Adams DW, Wu LJ, Errington J. 2016. A benzamide-dependent ftsZ mutant reveals residues crucial for Z-ring assembly. Mol Microbiol 99: 1028–1042.

Addinall SG, Holland B. 2002. The tubulin ancestor, FtsZ, draughtsman, designer and driving force for bacterial cytokinesis. J Mol Biol 318: 219–236.

Addinall SG, Lutkenhaus J. 1996. FtsZ-spirals and -arcs determine the shape of the invaginating septa in some mutants of Escherichia coli. Mol Microbiol 22: 231–237.

Agard D. 1984. Optical sectioning microscopy: cellular architecture in three dimensions. Annu Rev Biophys Biomol Struct 13: 191–219.

Anderson DE, Gueiros-Filho FJ, Erickson HP. 2004. Assembly dynamics of FtsZ rings in Bacillus subtilis and Escherichia coli and effects of FtsZ-regulating proteins. J Bacteriol 186: 5775–5781.

Anderson DE, Kim MB, Moore JT, O’Brien TE, Sorto NA, Grove CI, Lackner LL, Ames JB, Shaw JT. 2012. Comparison of small molecule inhibitors of the bacterial cell division protein FtsZ and identification of a reliable cross-species inhibitor. ACS Chem Biol 7: 1918–1928.

Andreu JM, Huecas S, Araújo-Bazán L, Vázquez-Villa H, Martín-Fontecha M. 2022. The search for antibacterial inhibitors targeting cell division protein ftsz at its nucleotide and allosteric binding sites. Biomedicines 10.

Andreu JM, Schaffner-Barbero C, Huecas S, Alonso D, Lopez-Rodriguez ML, Ruiz-Avila LB, Núñez-Ramírez R, Llorca O, Martín-Galiano AJ. 2010. The antibacterial cell division inhibitor PC190723 is an FtsZ polymer-stabilizing agent that induces filament assembly and condensation. J Biol Chem 285: 14239–14246.

Araújo-Bazán L, Huecas S, Valle J, Andreu D, Andreu JM. 2019. Synthetic developmental regulator MciZ targets FtsZ across Bacillus species and inhibits bacterial division. Mol Microbiol 111: 965–980.

Araújo-Bazán L, Ruiz-Avila LB, Andreu D, Huecas S, Andreu JM. 2016. Cytological profile of antibacterial ftsz inhibitors and synthetic peptide mciz. Front Microbiol 7: 1558.

Bähler J, Wu J-Q, Longtine MS, Shah NG, Mckenzie III A, Steever AB, Wach A, Philippsen P, Pringle JR. 1998. Heterologous modules for efficient and versatile PCR-based gene targeting in *Schizosaccharomyces pombe*. Yeast 14: 943–951.

Balzi E, Goffeau A. 1991. Multiple or pleiotropic drug resistance in yeast. Biochimica et Biophysica Acta (BBA) - General Subjects 1073: 241–252.

Barrows JM, Goley ED. 2021. FtsZ dynamics in bacterial division: What, how, and why? Curr Opin Cell Biol 68: 163–172.

Basi G, Schmid E, Maundrell K. 1993. TATA box mutations in the Schizosaccharomyces pombe nmt1 promoter affect transcription efficiency but not the transcription start point or thiamine repressibility. Gene 123: 131–136.

Benko Z, Liang D, Li G, Elder RT, Sarkar A, Takayama J, Ghosh AK, Zhao RY. 2017. A fission yeast cell-based system for multidrug resistant HIV-1 proteases. Cell Biosci 7: 5.

Benko Z, Zhang J, Zhao RY. 2019. Development of A Fission Yeast Cell-Based Platform for High Throughput Screening of HIV-1 Protease Inhibitors. Curr HIV Res 17: 429–440.

Beuria TK, Santra MK, Panda D. 2005. Sanguinarine blocks cytokinesis in bacteria by inhibiting FtsZ assembly and bundling. Biochemistry 44: 16584–16593.

Bi EF, Lutkenhaus J. 1991. FtsZ ring structure associated with division in Escherichia coli. Nature 354: 161–164.

Boberek JM, Stach J, Good L. 2010. Genetic evidence for inhibition of bacterial division protein FtsZ by berberine. PLoS ONE 5: e13745.

Bramhill D, Thompson CM. 1994. GTP-dependent polymerization of Escherichia coli FtsZ protein to form tubules. Proc Natl Acad Sci USA 91: 5813–5817.

Butler MS, Paterson DL. 2020. Antibiotics in the clinical pipeline in October 2019. J Antibiot 73: 329–364.

Carro L. 2019. Recent Progress in the Development of Small-Molecule FtsZ Inhibitors as Chemical Tools for the Development of Novel Antibiotics. Antibiotics (Basel) 8.

Casiraghi A, Suigo L, Valoti E, Straniero V. 2020. Targeting Bacterial Cell Division: A Binding Site-Centered Approach to the Most Promising Inhibitors of the Essential Protein FtsZ. Antibiotics (Basel) 9.

Cernáková M, Kostálová D. 2002. Antimicrobial activity of berberine--a constituent of Mahonia aquifolium. Folia Microbiol (Praha) 47: 375–378.

Chai WC, Whittall JJ, Song D, Polyak SW, Ogunniyi AD, Wang Y, Bi F, Ma S, Semple SJ, Venter H. 2020. Antimicrobial action and reversal of resistance in MRSA by difluorobenzamide derivatives targeted at FtsZ. Antibiotics (Basel) 9.

Chan F-Y, Sun N, Neves MAC, Lam PC-H, Chung W-H, Wong L-K, Chow H-Y, Ma D-L, Chan P-H, Leung Y-C, et al. 2013. Identification of a new class of FtsZ inhibitors by structure-based design and in vitro screening. J Chem Inf Model 53: 2131–2140.

Chu M, Zhang M-B, Liu Y-C, Kang J-R, Chu Z-Y, Yin K-L, Ding L-Y, Ding R, Xiao R-X, Yin Y-N, et al. 2016. Role of Berberine in the Treatment of Methicillin-Resistant Staphylococcus aureus Infections. Sci Rep 6: 24748.

Claridge-Chang A, Assam PN. 2016. Estimation statistics should replace significance testing. Nat Methods 13: 108–109.

Cumming G, Fidler F, Vaux DL. 2007. Error bars in experimental biology. J Cell Biol 177: 7–11.

Di Somma A, Avitabile C, Cirillo A, Moretta A, Merlino A, Paduano L, Duilio A, Romanelli A. 2020. The antimicrobial peptide Temporin L impairs E. coli cell division by interacting with FtsZ and the divisome complex. Biochim Biophys Acta Gen Subj 1864: 129606.

Domadia P, Swarup S, Bhunia A, Sivaraman J, Dasgupta D. 2007. Inhibition of bacterial cell division protein FtsZ by cinnamaldehyde. Biochem Pharmacol 74: 831–840.

Domadia PN, Bhunia A, Sivaraman J, Swarup S, Dasgupta D. 2008. Berberine targets assembly of Escherichia coli cell division protein FtsZ. Biochemistry 47: 3225–3234.

Elsen NL, Lu J, Parthasarathy G, Reid JC, Sharma S, Soisson SM, Lumb KJ. 2012. Mechanism of action of the cell-division inhibitor PC190723: modulation of FtsZ assembly cooperativity. J Am Chem Soc 134: 12342–12345.

Erickson HP, Anderson DE, Osawa M. 2010. FtsZ in bacterial cytokinesis: cytoskeleton and force generator all in one. Microbiol Mol Biol Rev 74: 504–528.

Erickson HP. 1995. FtsZ, a prokaryotic homolog of tubulin? Cell 80: 367–370.

Fujita J, Maeda Y, Mizohata E, Inoue T, Kaul M, Parhi AK, LaVoie EJ, Pilch DS, Matsumura H. 2017. Structural Flexibility of an Inhibitor Overcomes Drug Resistance Mutations in Staphylococcus aureus FtsZ. ACS Chem Biol 12: 1947–1955.

Gitai Z, Dye NA, Reisenauer A, Wachi M, Shapiro L. 2005. MreB actin-mediated segregation of a specific region of a bacterial chromosome. Cell 120: 329–341.

Haydon DJ, Stokes NR, Ure R, Galbraith G, Bennett JM, Brown DR, Baker PJ, Barynin VV, Rice DW, Sedelnikova SE, et al. 2008. An inhibitor of FtsZ with potent and selective anti-staphylococcal activity. Science 321: 1673–1675.

Henty-Ridilla JL, Li J, Day B, Staiger CJ. 2014. ACTIN DEPOLYMERIZING FACTOR4 regulates actin dynamics during innate immune signaling in Arabidopsis. Plant Cell 26: 340–352.

Hernández-Rocamora VM, Alfonso C, Margolin W, Zorrilla S, Rivas G. 2015. Evidence that bacteriophage λ kil peptide inhibits bacterial cell division by disrupting ftsz protofilaments and sequestering protein subunits. J Biol Chem 290: 20325–20335.

Higaki T. 2017. Quantitative evaluation of cytoskeletal organizations by microscopic image analysis. PLANT MORPHOL 29: 15–21.

Hou S, Wieczorek SA, Kaminski TS, Ziebacz N, Tabaka M, Sorto NA, Foss MH, Shaw JT, Thanbichler M, Weibel DB, et al. 2012. Characterization of Caulobacter crescentus FtsZ protein using dynamic light scattering. J Biol Chem 287: 23878–23886.

Ho J, Tumkaya T, Aryal S, Choi H, Claridge-Chang A. 2019. Moving beyond P values: data analysis with estimation graphics. Nat Methods 16: 565–566.

Huang K-H, Durand-Heredia J, Janakiraman A. 2013. FtsZ ring stability: of bundles, tubules, crosslinks, and curves. J Bacteriol 195: 1859–1868.

Huecas S, Ramírez-Aportela E, Vergoñós A, Núñez-Ramírez R, Llorca O, Díaz JF, Juan-Rodríguez D, Oliva MA, Castellen P, Andreu JM. 2017. Self-Organization of FtsZ Polymers in Solution Reveals Spacer Role of the Disordered C-Terminal Tail. Biophys J 113: 1831–1844.

Hurley KA, Santos TM, Nepomuceno GM, Huynh V, Shaw JT, Weibel DB. 2016. Targeting the bacterial division protein FtsZ. J Med Chem 59: 6975–6998.

Irwin JJ, Duan D, Torosyan H, Doak AK, Ziebart KT, Sterling T, Tumanian G, Shoichet BK. 2015. An aggregation advisor for ligand discovery. J Med Chem 58: 7076–7087.

Iwai N, Nagai K, Wachi M. 2002. Novel S-benzylisothiourea compound that induces spherical cells in Escherichia coli probably by acting on a rod-shape-determining protein(s) other than penicillin-binding protein 2. Biosci Biotechnol Biochem 66: 2658–2662.

Kaul M, Mark L, Zhang Y, Parhi AK, LaVoie EJ, Pilch DS. 2013a. Pharmacokinetics and in vivo antistaphylococcal efficacy of TXY541, a 1-methylpiperidine-4-carboxamide prodrug of PC190723. Biochem Pharmacol 86: 1699–1707.

Kaul M, Parhi AK, Zhang Y, LaVoie EJ, Tuske S, Arnold E, Kerrigan JE, Pilch DS. 2012. A bactericidal guanidinomethyl biaryl that alters the dynamics of bacterial FtsZ polymerization. J Med Chem 55: 10160–10176.

Kaul M, Zhang Y, Parhi AK, Lavoie EJ, Pilch DS. 2014. Inhibition of RND-type efflux pumps confers the FtsZ-directed prodrug TXY436 with activity against Gram-negative bacteria. Biochem Pharmacol 89: 321–328.

Kaul M, Zhang Y, Parhi AK, Lavoie EJ, Tuske S, Arnold E, Kerrigan JE, Pilch DS. 2013b. Enterococcal and streptococcal resistance to PC190723 and related compounds: molecular insights from a FtsZ mutational analysis. Biochimie 95: 1880–1887.

Kelley C, Zhang Y, Parhi A, Kaul M, Pilch DS, LaVoie EJ. 2012. 3-Phenyl substituted 6,7-dimethoxyisoquinoline derivatives as FtsZ-targeting antibacterial agents. Bioorg Med Chem 20: 7012–7029.

Khare S, Hsin J, Sorto NA, Nepomuceno GM, Shaw JT, Shi H, Huang KC. 2019. FtsZ- Independent Mechanism of Division Inhibition by the Small Molecule PC190723 in Escherichia coli. Adv Biosys 3: e1900021.

Kumar K, Awasthi D, Lee S-Y, Zanardi I, Ruzsicska B, Knudson S, Tonge PJ, Slayden RA, Ojima I. 2011. Novel trisubstituted benzimidazoles, targeting Mtb FtsZ, as a new class of antitubercular agents. J Med Chem 54: 374–381.

Kusuma KD, Payne M, Ung AT, Bottomley AL, Harry EJ. 2019. FtsZ as an antibacterial target: status and guidelines for progressing this avenue. ACS Infect Dis 5: 1279–1294.

Läppchen T, Pinas VA, Hartog AF, Koomen G-J, Schaffner-Barbero C, Andreu JM, Trambaiolo D, Löwe J, Juhem A, Popov AV, et al. 2008. Probing FtsZ and tubulin with C8-substituted GTP analogs reveals differences in their nucleotide binding sites. Chem Biol 15: 189–199.

Liu F, Venter H, Bi F, Semple SJ, Liu J, Jin C, Ma S. 2017. Synthesis and antibacterial activity of 5-methylphenanthridium derivatives as FtsZ inhibitors. Bioorg Med Chem Lett 27: 3399–3402.

Li X, Sheng J, Huang G, Ma R, Yin F, Song D, Zhao C, Ma S. 2015. Design, synthesis and antibacterial activity of cinnamaldehyde derivatives as inhibitors of the bacterial cell division protein FtsZ. Eur J Med Chem 97: 32–41.

Lopus M, Panda D. 2006. The benzophenanthridine alkaloid sanguinarine perturbs microtubule assembly dynamics through tubulin binding. A possible mechanism for its antiproliferative activity. FEBS J 273: 2139–2150.

Lord SJ, Velle KB, Mullins RD, Fritz-Laylin LK. 2020. SuperPlots: Communicating reproducibility and variability in cell biology. J Cell Biol 219.

Löwe J, Amos LA. 1998. Crystal structure of the bacterial cell-division protein FtsZ. Nature 391: 203–206.

Lui HK, Gao W, Cheung KC, Jin WB, Sun N, Kan JWY, Wong ILK, Chiou J, Lin D, Chan EWC, et al. 2019. Boosting the efficacy of anti-MRSA β-lactam antibiotics via an easily accessible, non-cytotoxic and orally bioavailable FtsZ inhibitor. Eur J Med Chem 163: 95–115.

Mahone CR, Goley ED. 2020. Bacterial cell division at a glance. J Cell Sci 133.

Margalit DN, Romberg L, Mets RB, Hebert AM, Mitchison TJ, Kirschner MW, RayChaudhuri D. 2004. Targeting cell division: small-molecule inhibitors of FtsZ GTPase perturb cytokinetic ring assembly and induce bacterial lethality. Proc Natl Acad Sci USA 101: 11821–11826.

Margolin W. 2005. FtsZ and the division of prokaryotic cells and organelles. Nat Rev Mol Cell Biol 6: 862–871.

Martin JK, Sheehan JP, Bratton BP, Moore GM, Mateus A, Li SH-J, Kim H, Rabinowitz JD, Typas A, Savitski MM, et al. 2020. A Dual-Mechanism Antibiotic Kills Gram-Negative Bacteria and Avoids Drug Resistance. Cell 181: 1518–1532.e14.

Matsui T, Yamane J, Mogi N, Yamaguchi H, Takemoto H, Yao M, Tanaka I. 2012a. Structural reorganization of the bacterial cell-division protein FtsZ from Staphylococcus aureus. Acta Crystallogr D Biol Crystallogr 68: 1175–1188.

Matsui T, Yamane J, Mogi N, Yamaguchi H, Takemoto H, Yao M, Tanaka I. 2012b. Structural reorganization of the bacterial cell-division protein FtsZ from Staphylococcus aureus. Acta Crystallogr D Biol Crystallogr 68: 1175–1188.

Ma X, Ehrhardt DW, Margolin W. 1996. Colocalization of cell division proteins FtsZ and FtsA to cytoskeletal structures in living Escherichia coli cells by using green fluorescent protein. Proc Natl Acad Sci USA 93: 12998–13003.

Miguel A, Hsin J, Liu T, Tang G, Altman RB, Huang KC. 2015. Variations in the binding pocket of an inhibitor of the bacterial division protein FtsZ across genotypes and species. PLoS Comput Biol 11: e1004117.

Mikuni S, Kodama K, Sasaki A, Kohira N, Maki H, Munetomo M, Maenaka K, Kinjo M. 2015. Screening for FtsZ Dimerization Inhibitors Using Fluorescence Cross-Correlation Spectroscopy and Surface Resonance Plasmon Analysis. PLoS ONE 10: e0130933.

Monahan LG, Robinson A, Harry EJ. 2009. Lateral FtsZ association and the assembly of the cytokinetic Z ring in bacteria. Mol Microbiol 74: 1004–1017.

Monterroso B, Alfonso C, Zorrilla S, Rivas G. 2013. Combined analytical ultracentrifugation, light scattering and fluorescence spectroscopy studies on the functional associations of the bacterial division FtsZ protein. Methods 59: 349–362.

Moreno MB, Durán A, Carlos Ribas J. 2000. A family of multifunctional thiamine-repressible expression vectors for fission yeast. Yeast 16: 861–872.

Mukherjee A, Cao C, Lutkenhaus J. 1998. Inhibition of FtsZ polymerization by SulA, an inhibitor of septation in Escherichia coli. Proc Natl Acad Sci USA 95: 2885–2890.

Mukherjee A, Lutkenhaus J. 1999. Analysis of FtsZ assembly by light scattering and determination of the role of divalent metal cations. J Bacteriol 181: 823–832.

Nepomuceno GM, Chan KM, Huynh V, Martin KS, Moore JT, O’Brien TE, Pollo LAE, Sarabia FJ, Tadeus C, Yao Z, et al. 2015. Synthesis and evaluation of quinazolines as inhibitors of the bacterial cell division protein ftsz. ACS Med Chem Lett 6: 308–312.

Nogales E, Downing KH, Amos LA, Löwe J. 1998. Tubulin and FtsZ form a distinct family of GTPases. Nat Struct Biol 5: 451–458.

Nonejuie P, Burkart M, Pogliano K, Pogliano J. 2013. Bacterial cytological profiling rapidly identifies the cellular pathways targeted by antibacterial molecules. Proc Natl Acad Sci USA 110: 16169–16174.

Nova E, Montecinos F, Brunet JE, Lagos R, Monasterio O. 2007. 4’,6-Diamidino-2-phenylindole (DAPI) induces bundling of Escherichia coli FtsZ polymers inhibiting the GTPase activity. Arch Biochem Biophys 465: 315–319.

Obiang-Obounou BW, Kang O-H, Choi J-G, Keum J-H, Kim S-B, Mun S-H, Shin D-W, Kim KW, Park C-B, Kim Y-G, et al. 2011. The mechanism of action of sanguinarine against methicillin-resistant Staphylococcus aureus. J Toxicol Sci 36: 277–283.

Pande V, Mitra N, Bagde SR, Srinivasan R, Gayathri P. 2022. Filament organization of the bacterial actin MreB is dependent on the nucleotide state. J Cell Biol 221.

Park B, Awasthi D, Chowdhury SR, Melief EH, Kumar K, Knudson SE, Slayden RA, Ojima I. 2014. Design, synthesis and evaluation of novel 2,5,6-trisubstituted benzimidazoles targeting FtsZ as antitubercular agents. Bioorg Med Chem 22: 2602–2612.

Pérez P, Ribas JC. 2004. Cell wall analysis. Methods 33: 245–251.

Pettersen EF, Goddard TD, Huang CC, Couch GS, Greenblatt DM, Meng EC, Ferrin TE. 2004. UCSF Chimera—a visualization system for exploratory research and analysis. J Comput Chem 25: 1605–1612.

Piddock LJV. 2006. Clinically relevant chromosomally encoded multidrug resistance efflux pumps in bacteria. Clin Microbiol Rev 19: 382–402.

Pierpaoli E, Cirioni O, Simonetti O, Orlando F, Giacometti A, Lombardi P, Provinciali M. 2021. Potential application of berberine in the treatment of Escherichia coli sepsis. Nat Prod Res 35: 4779–4784.

Pollard DA, Pollard TD, Pollard KS. 2019. Empowering statistical methods for cellular and molecular biologists. Mol Biol Cell 30: 1359–1368.

Popp D, Iwasa M, Narita A, Erickson HP, Maéda Y. 2009. FtsZ condensates: an in vitro electron microscopy study. Biopolymers 91: 340–350.

Pradhan P, Margolin W, Beuria TK. 2021. Targeting the achilles heel of ftsz: the interdomain cleft. Front Microbiol 12: 732796.

Raghav D, Ashraf SM, Mohan L, Rathinasamy K. 2017. Berberine Induces Toxicity in HeLa Cells through Perturbation of Microtubule Polymerization by Binding to Tubulin at a Unique Site. Biochemistry 56: 2594–2611.

Rai D, Singh JK, Roy N, Panda D. 2008. Curcumin inhibits FtsZ assembly: an attractive mechanism for its antibacterial activity. Biochem J 410: 147–155.

RayChaudhuri D, Park JT. 1992. Escherichia coli cell-division gene ftsZ encodes a novel GTP-binding protein. Nature 359: 251–254.

Reija B, Monterroso B, Jiménez M, Vicente M, Rivas G, Zorrilla S. 2011. Development of a homogeneous fluorescence anisotropy assay to monitor and measure FtsZ assembly in solution. Anal Biochem 418: 89–96.

Rivas G, Vogel SK, Schwille P. 2014. Reconstitution of cytoskeletal protein assemblies for large-scale membrane transformation. Curr Opin Chem Biol 22: 18–26.

Rosado-Lugo JD, Sun Y, Banerjee A, Cao Y, Datta P, Zhang Y, Yuan Y, Parhi AK. 2022. Evaluation of 2,6-difluoro-3-(oxazol-2-ylmethoxy)benzamide chemotypes as Gram-negative FtsZ inhibitors. J Antibiot 75: 385–395.

Ruiz-Avila LB, Huecas S, Artola M, Vergoñós A, Ramírez-Aportela E, Cercenado E, Barasoain I, Vázquez-Villa H, Martín-Fontecha M, Chacón P, et al. 2013. Synthetic inhibitors of bacterial cell division targeting the GTP-binding site of FtsZ. ACS Chem Biol 8: 2072–2083.

Schaffner-Barbero C, Martín-Fontecha M, Chacón P, Andreu JM. 2012. Targeting the assembly of bacterial cell division protein FtsZ with small molecules. ACS Chem Biol 7: 269–277.

Schindelin J, Arganda-Carreras I, Frise E, Kaynig V, Longair M, Pietzsch T, Preibisch S, Rueden C, Saalfeld S, Schmid B, et al. 2012. Fiji: an open-source platform for biological-image analysis. Nat Methods 9: 676–682.

Sievers F, Wilm A, Dineen D, Gibson TJ, Karplus K, Li W, Lopez R, McWilliam H, Remmert M, Söding J, et al. 2011. Fast, scalable generation of high-quality protein multiple sequence alignments using Clustal Omega. Mol Syst Biol 7: 539.

Silber N, Matos de Opitz CL, Mayer C, Sass P. 2020. Cell division protein FtsZ: from structure and mechanism to antibiotic target. Future Microbiol 15: 801–831.

Spahn C, Gómez-de-Mariscal E, Laine RF, Pereira PM, von Chamier L, Conduit M, Pinho MG, Jacquemet G, Holden S, Heilemann M, et al. 2022. DeepBacs for multi-task bacterial image analysis using open-source deep learning approaches. Commun Biol 5: 688.

Specht M, Dempwolff F, Schätzle S, Thomann R, Waidner B. 2013. Localization of FtsZ in Helicobacter pylori and consequences for cell division. J Bacteriol 195: 1411–1420.

Srinivasan R, Mishra M, Murata-Hori M, Balasubramanian MK. 2007. Filament formation of the Escherichia coli actin-related protein, MreB, in fission yeast. Curr Biol 17: 266–272.

Srinivasan R, Mishra M, Wu L, Yin Z, Balasubramanian MK. 2008. The bacterial cell division protein FtsZ assembles into cytoplasmic rings in fission yeast. Genes Dev 22: 1741–1746.

Stokes NR, Baker N, Bennett JM, Berry J, Collins I, Czaplewski LG, Logan A, Macdonald R, Macleod L, Peasley H, et al. 2013. An improved small-molecule inhibitor of FtsZ with superior in vitro potency, drug-like properties, and in vivo efficacy. Antimicrob Agents Chemother 57: 317–325.

Strahl H, Hamoen LW. 2010. Membrane potential is important for bacterial cell division. Proc Natl Acad Sci USA 107: 12281–12286.

Straniero V, Sebastián-Pérez V, Hrast M, Zanotto C, Casiraghi A, Suigo L, Zdovc I, Radaelli A, De Giuli Morghen C, Valoti E. 2020a. Benzodioxane-Benzamides as Antibacterial Agents: Computational and SAR Studies to Evaluate the Influence of the 7-Substitution in FtsZ Interaction. ChemMedChem 15: 195–209.

Straniero V, Suigo L, Casiraghi A, Sebastián-Pérez V, Hrast M, Zanotto C, Zdovc I, De Giuli Morghen C, Radaelli A, Valoti E. 2020b. Benzamide derivatives targeting the cell division protein ftsz: modifications of the linker and the benzodioxane scaffold and their effects on antimicrobial activity. Antibiotics (Basel) 9.

Straniero V, Zanotto C, Straniero L, Casiraghi A, Duga S, Radaelli A, De Giuli Morghen C, Valoti E. 2017. 2,6-Difluorobenzamide Inhibitors of Bacterial Cell Division Protein FtsZ: Design, Synthesis, and Structure-Activity Relationships. ChemMedChem 12: 1303–1318.

Sun N, Chan F-Y, Lu Y-J, Neves MAC, Lui H-K, Wang Y, Chow K-Y, Chan K-F, Yan S-C, Leung Y-C, et al. 2014. Rational design of berberine-based FtsZ inhibitors with broad-spectrum antibacterial activity. PLoS ONE 9: e97514.

Sun N, Lu Y-J, Chan F-Y, Du R-L, Zheng Y-Y, Zhang K, So L-Y, Abagyan R, Zhuo C, Leung Y-C, et al. 2017. A thiazole orange derivative targeting the bacterial protein ftsz shows potent antibacterial activity. Front Microbiol 8: 855.

Tan CM, Therien AG, Lu J, Lee SH, Caron A, Gill CJ, Lebeau-Jacob C, Benton-Perdomo L, Monteiro JM, Pereira PM, et al. 2012. Restoring methicillin-resistant Staphylococcus aureus susceptibility to β-lactam antibiotics. Sci Transl Med 4: 126ra35.

TAXIS Pharmaceuticals. 2020. TAXIS secures NIH Funding – TAXIS Pharmaceutical. https://www.taxispharma.com/news/taxis-secures-nih-funding/ (Accessed February 8, 2022).

TerBush AD, MacCready JS, Chen C, Ducat DC, Osteryoung KW. 2018. Conserved dynamics of chloroplast cytoskeletal ftsz proteins across photosynthetic lineages. Plant Physiol 176: 295–306.

TerBush AD, Osteryoung KW. 2012. Distinct functions of chloroplast FtsZ1 and FtsZ2 in Z-ring structure and remodeling. J Cell Biol 199: 623–637.

Tripathy S, Sahu SK. 2019. FtsZ inhibitors as a new genera of antibacterial agents. Bioorg Chem 91: 103169.

van den Ent F, Löwe J. 2006. RF cloning: a restriction-free method for inserting target genes into plasmids. J Biochem Biophys Methods 67: 67–74.

Vaughan S, Wickstead B, Gull K, Addinall SG. 2004. Molecular evolution of FtsZ protein sequences encoded within the genomes of archaea, bacteria, and eukaryota. J Mol Evol 58: 19–29.

Vaux DL. 2014. Basic statistics in cell biology. Annu Rev Cell Dev Biol 30: 23–37.

Vollmer W. 2006. The prokaryotic cytoskeleton: a putative target for inhibitors and antibiotics? Appl Microbiol Biotechnol 73: 37–47.

von Chamier L, Laine RF, Jukkala J, Spahn C, Krentzel D, Nehme E, Lerche M, Hernández-Pérez S, Mattila PK, Karinou E, et al. 2021. Democratising deep learning for microscopy with ZeroCostDL4Mic. Nat Commun 12: 2276.

Wang J, Galgoci A, Kodali S, Herath KB, Jayasuriya H, Dorso K, Vicente F, González A, Cully D, Bramhill D, et al. 2003. Discovery of a small molecule that inhibits cell division by blocking FtsZ, a novel therapeutic target of antibiotics. J Biol Chem 278: 44424–44428.

Wang J, Ma W, Fang Y, Liang H, Yang H, Wang Y, Dong X, Zhan Y, Wang X. 2021. Core Oligosaccharide Portion of Lipopolysaccharide Plays Important Roles in Multiple Antibiotic Resistance in Escherichia coli. Antimicrob Agents Chemother 65: e0034121.

Wang X, Tanaka M, Krstin S, Peixoto HS, Wink M. 2016. The Interference of Selected Cytotoxic Alkaloids with the Cytoskeleton: An Insight into Their Modes of Action. Molecules 21.

Wang Y, Yan M, Ma R, Ma S. 2015. Synthesis and antibacterial activity of novel 4-bromo-1H-indazole derivatives as FtsZ inhibitors. Arch Pharm (Weinheim) 348: 266–274.

Waterhouse A, Bertoni M, Bienert S, Studer G, Tauriello G, Gumienny R, Heer FT, de Beer TAP, Rempfer C, Bordoli L, et al. 2018. SWISS-MODEL: homology modelling of protein structures and complexes. Nucleic Acids Res 46: W296–W303.

Waterhouse AM, Procter JB, Martin DMA, Clamp M, Barton GJ. 2009. Jalview Version 2--a multiple sequence alignment editor and analysis workbench. Bioinformatics 25: 1189–1191.

Weiss DS, Chen JC, Ghigo JM, Boyd D, Beckwith J. 1999. Localization of FtsI (PBP3) to the septal ring requires its membrane anchor, the Z ring, FtsA, FtsQ, and FtsL. J Bacteriol 181: 508–520.

Weiss DS. 2004. Bacterial cell division and the septal ring. Mol Microbiol 54: 588–597.

Wolff J, Knipling L. 1993. Antimicrotubule properties of benzophenanthridine alkaloids. Biochemistry 32: 13334–13339.

Yoshida Y, Mogi Y, TerBush AD, Osteryoung KW. 2016. Chloroplast FtsZ assembles into a contractible ring via tubulin-like heteropolymerization. Nat Plants 2: 16095.

Zorrilla S, Monterroso B, Robles-Ramos M-Á, Margolin W, Rivas G. 2021. FtsZ interactions and biomolecular condensates as potential targets for new antibiotics. Antibiotics (Basel) 10.

