## Supplemental Figures for "A salt bridge-mediated resistance mechanism to FtsZ inhibitor PC190723 revealed by a cell-based screen"

### Supplementary Figures

### Figure S1

HpFtsZ-GFP

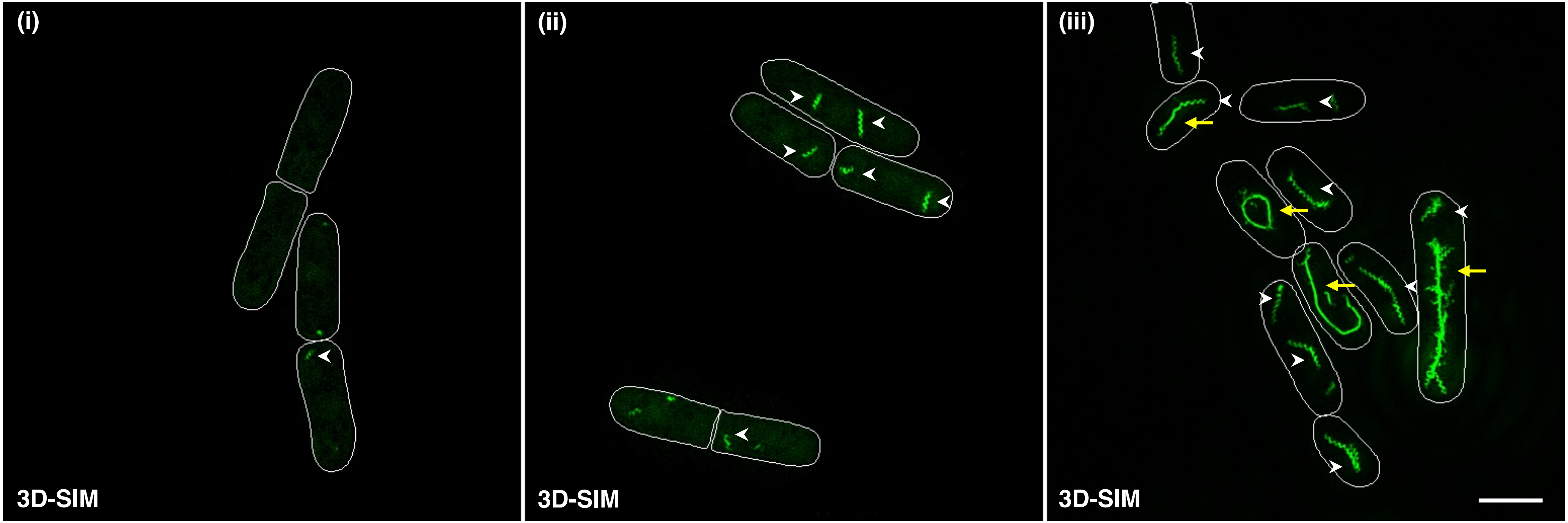


| 12 – 14 hours (MM – thiamine) | 16 – 18 hours (MM – thiamine) | 22 – 24 hours (MM – thiamine) |
|---|---|---|

| | **16 – 18 hours** | **22 – 24 hours** |
|---|---|---|
| **Ratio (Spiral / Linear filaments)** | 25.11 | 0.88 |

**Figure S2**

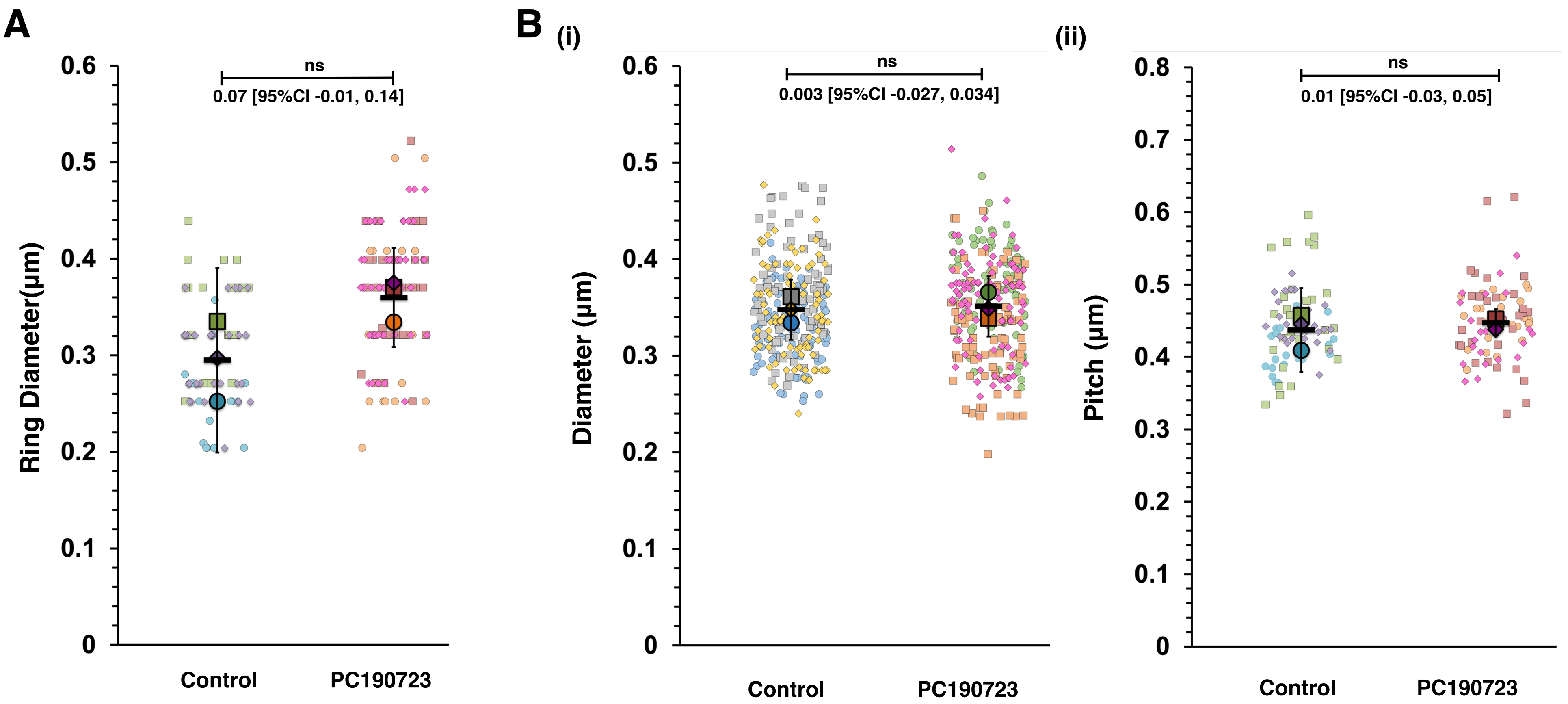

A
0.6
0.5
0.4
0.3
0.2
0.1
0
ns
0.07 [95%CI -0.01, 0.14]
Ring Diameter(µm)
Control
PC190723
B
(i)
0.6
0.5
0.4
0.3
0.2
0.1
0
ns
0.003 [95%CI -0.027, 0.034]
Diameter (µm)
Control
PC190723
(ii)
0.8
0.7
0.6
0.5
0.4
0.3
0.2
0.1
0
ns
0.01 [95%CI -0.03, 0.05]
Pitch (µm)
Control
PC190723

#### Figure S3

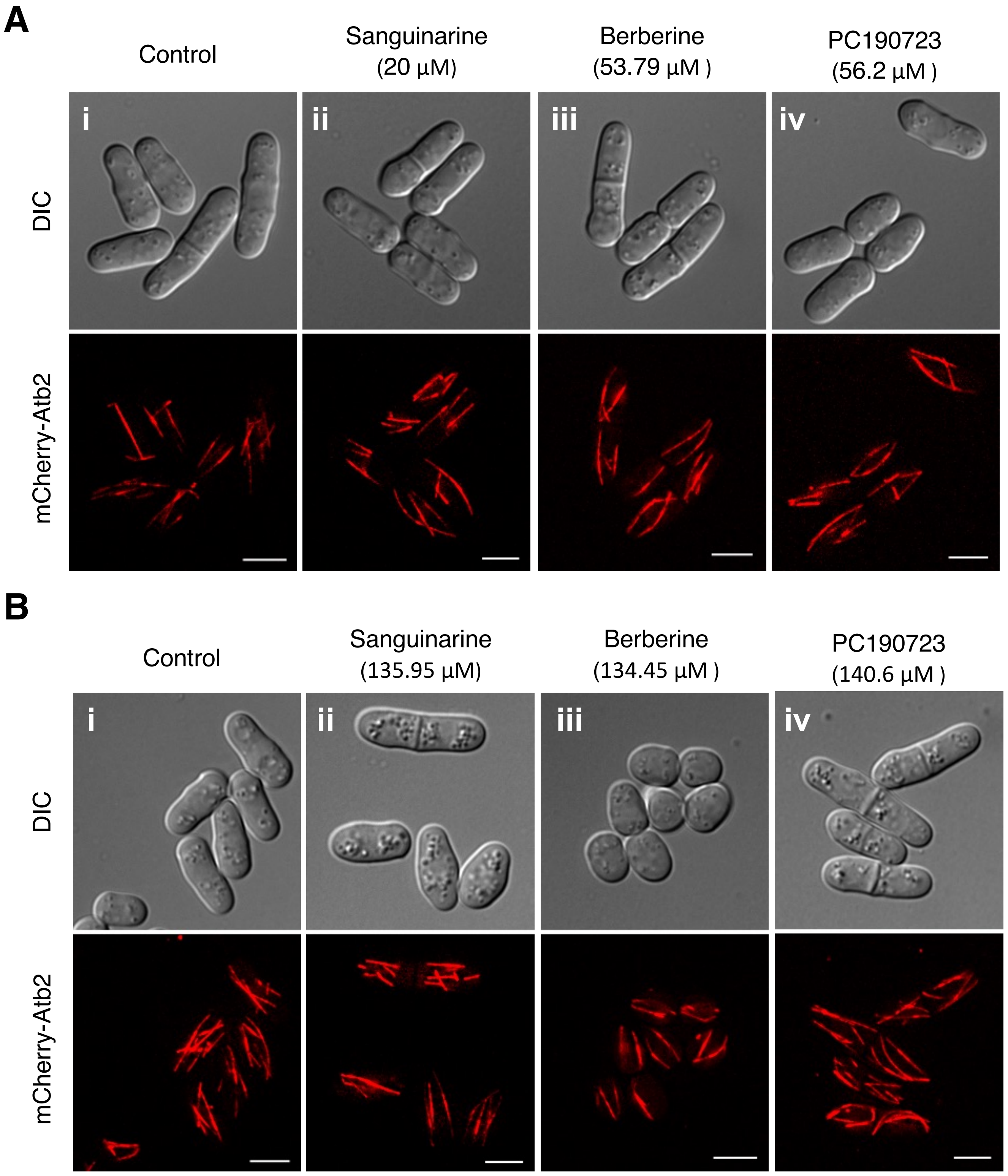
A
Control
Sanguinarine
(20 μM)
Berberine
(53.79 μM )
PC190723
(56.2 μM )
i
ii
iii
iv
DIC
mCherry-Atb2
B
Control
Sanguinarine
(135.95 μM)
Berberine
(134.45 μM )
PC190723
(140.6 μM )
i
ii
iii
iv
DIC
mCherry-Atb2

**Figure S3**

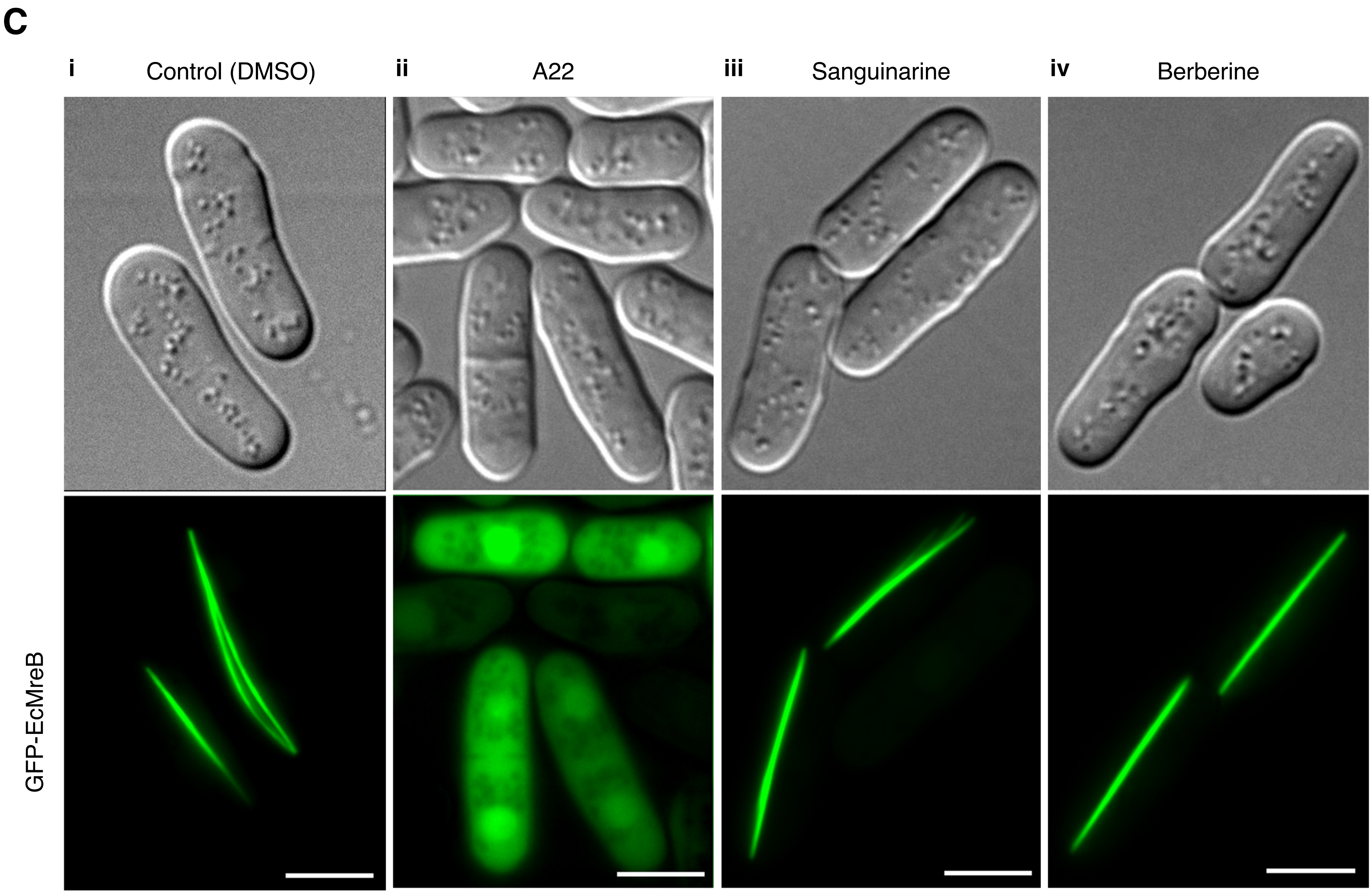
C
i
Control (DMSO)
ii
A22
iii
Sanguinarine
iv
Berberine
GFP-EcMreB

#### Figure S4

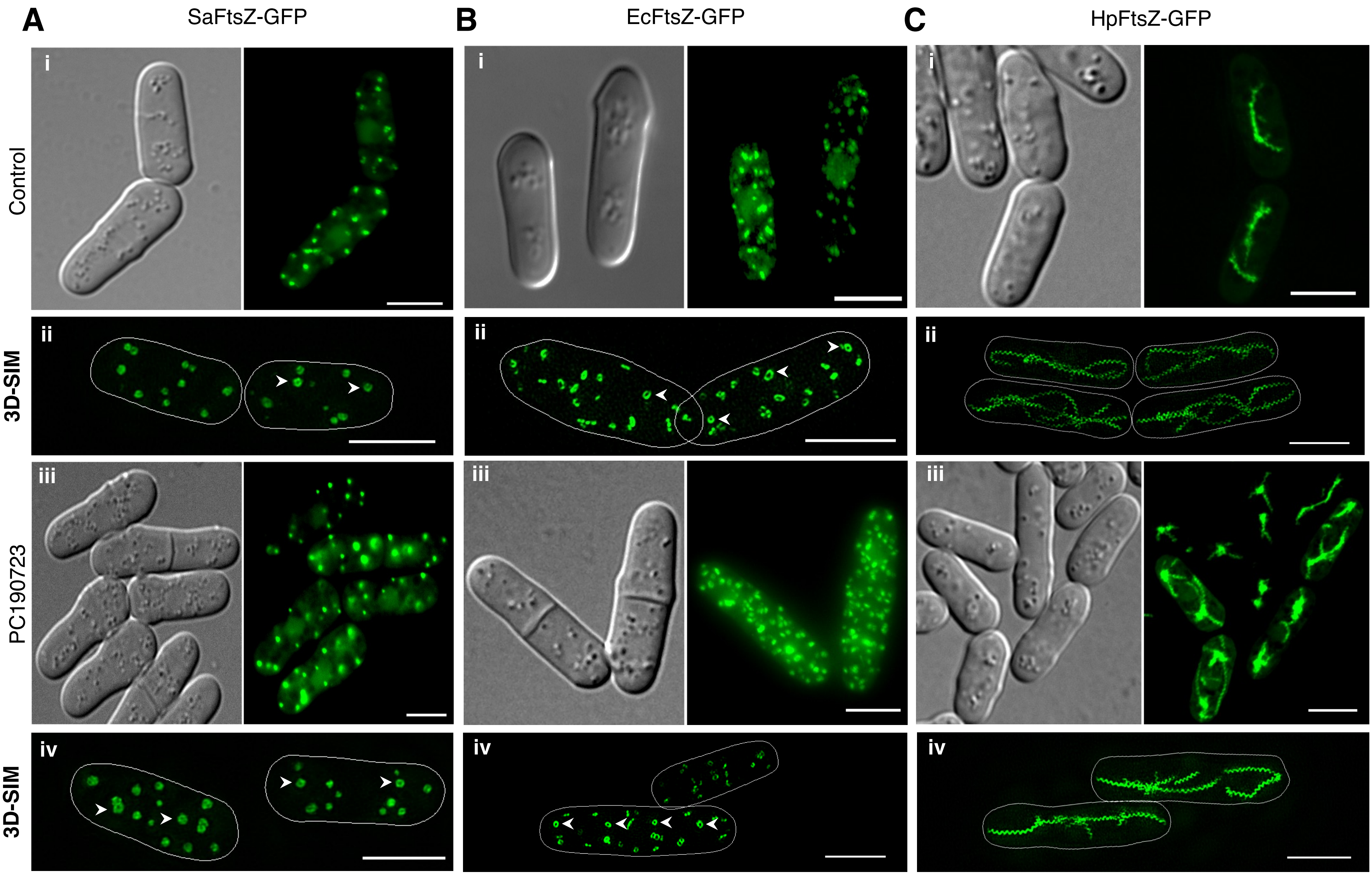

A
SaFtsZ-GFP
B
EcFtsZ-GFP
C
HpFtsZ-GFP
Control
3D-SIM
PC190723
3D-SIM
i
ii
iii
iv

### Figure S5

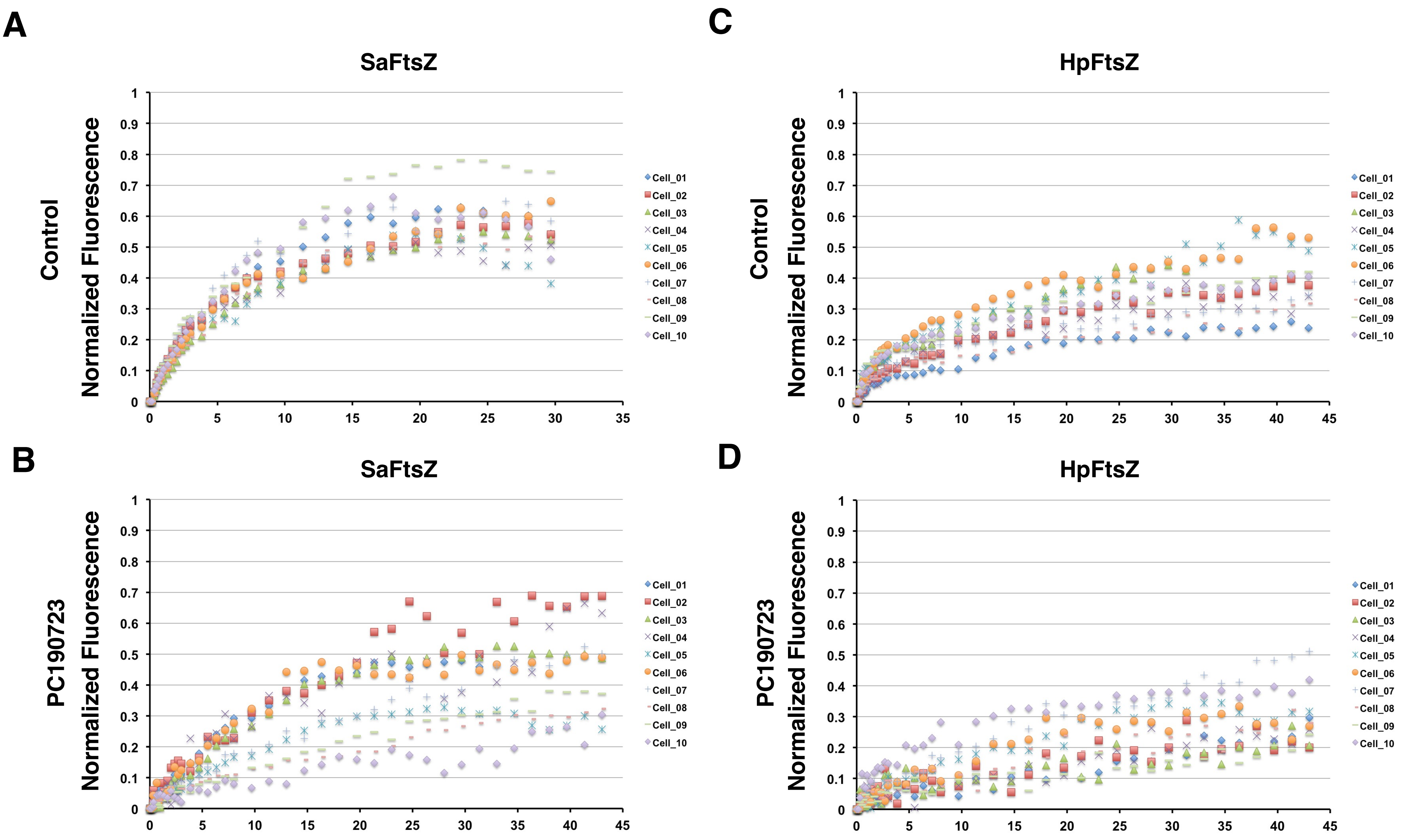
A
SaFtsZ
Control
Normalized Fluorescence
1
0.9
0.8
0.7
0.6
0.5
0.4
0.3
0.2
0.1
0
0
5
10
15
20
25
30
35
Cell_01
Cell_02
Cell_03
Cell_04
Cell_05
Cell_06
Cell_07
Cell_08
Cell_09
Cell_10
C
HpFtsZ
Control
Normalized Fluorescence
0
5
10
15
20
25
30
35
40
45
B
SaFtsZ
PC190723
Normalized Fluorescence
0
5
10
15
20
25
30
35
40
45
D
HpFtsZ
PC190723
Normalized Fluorescence
0
5
10
15
20
25
30
35
40
45

### Figure S6

A

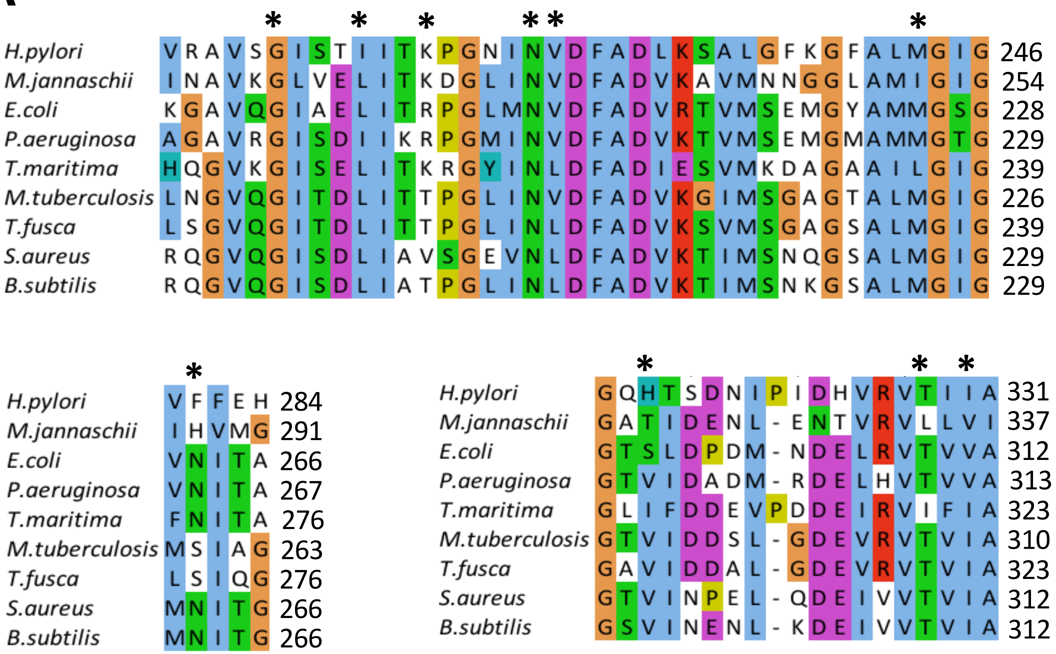


B

| **Residues interacting with PC190723 (SaFtsZ)** | **Corresponding residues in *Hp*FtsZ** |
|---|---|
| G196 | G213 |
| L200 | I217 |
| V203 | K220 (does not interact) |
| N208 | N225 |
| L209 | V226 |
| M226 | M243 |
| N263 | F281 (stacking) |
| V297 | H315 (does not interact) |
| T309 | T328 |
| I311 | I330 |

### Figure S7

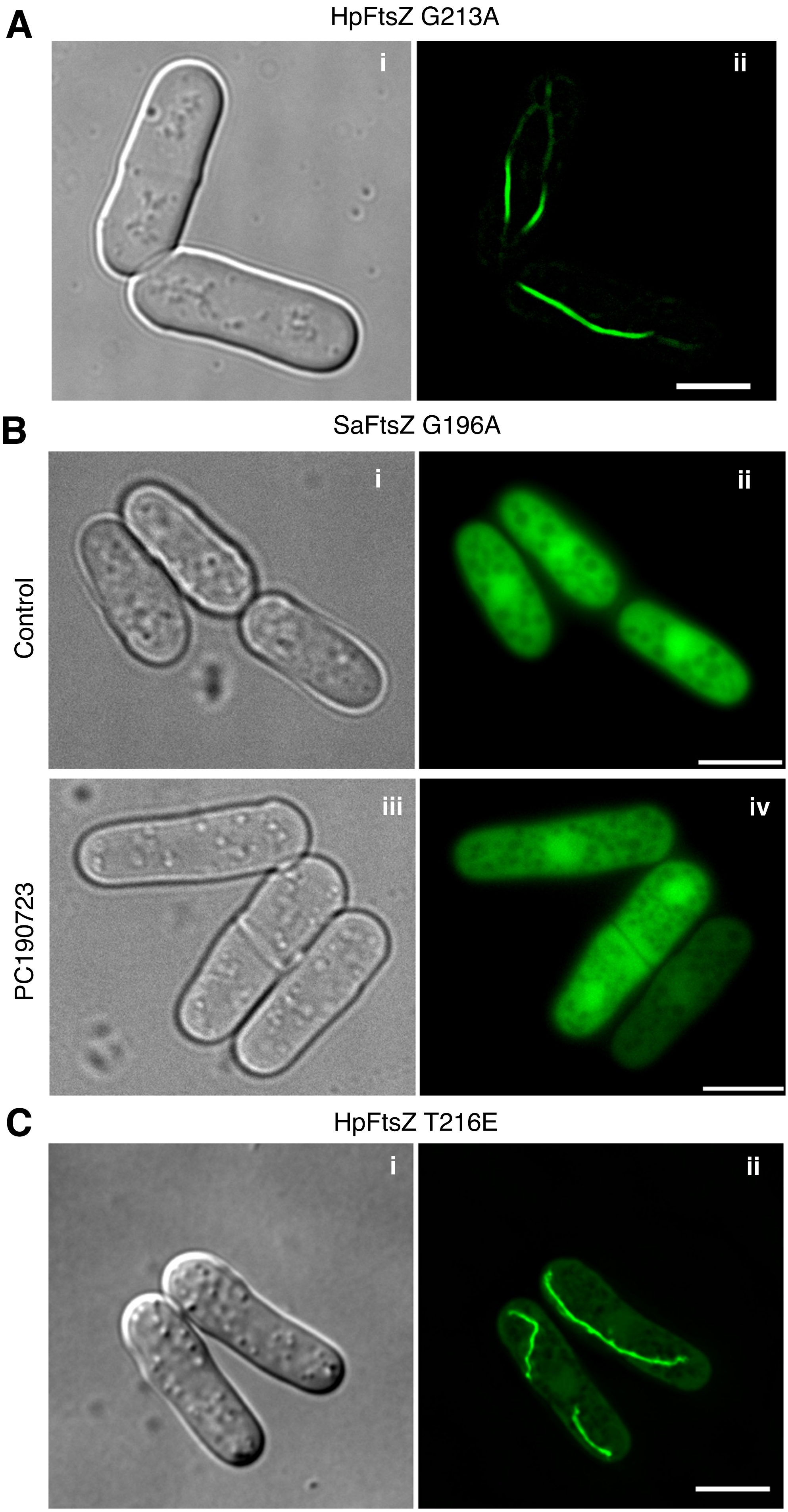
A
HpFtsZ G213A
i
ii
B
SaFtsZ G196A
i
ii
Control
iii
iv
PC190723
C
HpFtsZ T216E
i
ii

**Supplementary Figure Legends:**

**Figure S1. HpFtsZ assembles into spiral and linear filaments in fission yeast.** 3D-SIM images of HpFtsZ filaments in fission yeast cells after induction of expression in the absence of thiamine for **(i)** 12 – 14 hours, **(ii)** 16 – 18 hours and **(iii)** 22 – 24 hours. Spiral polymers of HpFtsZ start to appear after 16 – 18 hours of expression and possibly bundle to form linear cables at longer induction times. Scale bar represents 5 μm.

**Figure S2. Dimensions of ring-like structures of SaFtsZ and spiral polymers of HpFtsZ.** Superplots showing **(A)** diameter of the ring-like structures assembled by SaFtsZ, **(Bi)** diameter and **(Bii)** pitch of the spiral polymers assembled by HpFtsZ in the presence or absence of PC190723. Diameter and pitch were measured from 3D-SIM images of cells from three independent biological replicates (N = 3). In the case of SaFtsZ, the number of rings were measured for each replicate was > 19 and < 52. In the case of HpFtsZ, 12 to 18 filaments and 19 to 30 filaments were used to measure the diameter and pitch, respectively, of the spirals from each replicate. A two-sided unpaired Student's t-test was used to calculate the P-value, and ns indicates a P-value ≥ 0.05. The mean difference between the untreated and PC190723 treated populations is also indicated as effect size [95% CI, lower bound, upper bound]. Biological replicates are independent liquid cultures starting from a fresh patch from frozen glycerol stocks.

**Figure S3. Sanguinarine and berberine do not affect yeast microtubules or the polymerization of the *E. coli* actin homolog MreB.** Fission yeast cultures expressing mCherry-Atb2 fusions were grown at 30ºC in the absence of thiamine for 10 – 12 hours. **(A I and B i)** DMSO, **(A ii)** 20 μM sanguinarine, **(A iii)** 53.79 μM berberine and **(A iv)** 56.2 μM PC190723 were added to the cultures and further grown at 30ºC for another 10 – 12 hours prior to imaging. **(B ii)** 135.95 μM sanguinarine, **(B iii)** 134.35 μM berberine and **(B iv)** 140.6 μM PC190723 were added to the cultures and further grown at 30°C for another 10 – 12 hours prior to imaging. **(C)** Effect of sanguinarine and berberine on MreB. A22, a known inhibitor of MreB, was used as a positive control. **(i)** DMSO-treated cells showed a linear array of EcMreB polymers in fission yeast. **(ii)** MreB inhibitor A22 (73.64 μM) prevented polymerization of EcMreB. **(iii)** sanguinarine (20 μM) and **(iv)** berberine (53.79 μM) treated cells showed MreB bundles similar to control cells suggesting failure to inhibit MreB assembly. Scale bar represents 5 μm.

**Figure S4. PC190723 does not affect the FtsZ structures in fission yeast after the assembly of FtsZ.** Fission yeast cultures expressing different bacterial FtsZ-GFP fusions **(A)** EcFtsZ_GFP, **(B)** SaFtsZ_GFP and **(C)** HpFtsZ_GFP were mounted onto 1.6% agarose slides before imaging using an epifluorescence microscope. The addition of PC190723 (56.2 μM) after the appearance of FtsZ polymers shows no significant effect on FtsZ polymers **(A – C, panels iii)**. In control (untreated) cultures **(A – C, panels i)**, an equivalent amount of DMSO was added. 3D-SIM images showing the presence of ring-like (SaFtsZ and EcFtsZ) or spiral polymers (HpFtsZ) in the absence **(A – C, panels ii)** as well as the presence of PC190723 **(A – C, panels iv).** *S. pombe* cells expressing EcFtsZ, SaFtsZ or HpFtsZ were grown at 30°C in the absence of thiamine for 20 – 24 hours before DMSO or PC190723 was added to the cultures. The cultures were further incubated for 4 hours at 30°C and imaged. Scale bar represents 5 μm.

**Figure S5. PC190723 reduces the polymer turnover rates of SaFtsZ and HpFtsZ.** Graphs showing the fluorescence recovery after photobleaching (FRAP) of **(A & B)** SaFtsZ-GFP or **(C & D)** HpFtsZ-GFP assemblies in fission yeast. **(A & C)** Control (DMSO treated) cells and **(B & D)** PC190723 (56.2 μM) treated cells. Fluorescence intensities during recovery were quantified, and single normalized intensities were plotted against time.

**Figure S6.** **(A)** Multiple sequence alignment of FtsZs from different bacteria; stars (*) represent the residues in SaFtsZ, which interacts with PC190723. **(B)** Residues interacting with PC190723 in SaFtsZ and their corresponding residues in HpFtsZ.

**Figure S7**. Assembly of FtsZ mutants resistant to PC190723 in fission yeast. **(A)** HpFtsZ mutation G213A that makes it resistant to the drug PC190723 is capable of polymerization and assembly into filaments in fission yeast. **(B)** SaFtsZ$^{G196A}$ fails to assemble into spots or patches either in the absence of the drug (panel ii) or in the presence of 20 $\mu$g / ml PC190723 (panel iv). **(C)** HpFtsZ mutant T216E, which is resistant to the drug PC190723, is capable of polymerization and assembly into filaments in fission yeast. *S. pombe* cultures carrying pREP42-HpFtsZ$^{G213A}$ or pREP42-HpFtsZ$^{T216E}$ were grown for 10 – 12 hours and then sub-cultured into media containing PC190723 or an equivalent volume of DMSO. They were further grown at 30°C for 4 hours to allow the expression of the proteins and imaged under the epifluorescence microscope. Scale bar represents 5 µm.
