## Supplemental Tables for "A salt bridge-mediated resistance mechanism to FtsZ inhibitor PC190723 revealed by a cell-based screen"

**Supplementary Table – S1**

**List of strains and plasmids:**

| Strains/ Plasmids | Description | Reference |
|---|---|---|
| *Yeast Strains* | | |
| CCDY3 | MBY3497; CCDY346/pREP42- EcFtsZ-GFP | Srinivasan et al., 2008 |
| CCDY4 | MBY3532; CCDY346/pREP42- GFP-EcMreB | Srinivasan et al., 2007 |
| CCDY340 | CCDY346/pREP42- SaFtsZ-GFP | This work |
| CCDY341 | CCDY346/pREP42- HpFtsZ-GFP | This work |
| CCDY346 | MBY192; *Schizosaccharomyces pombe* [*ura4*-D18, *leu1*-32, h-] | Dr. Mithilesh Mishra (DBS, TIFR) |
| CCDY365 | MBY5856; *Schizosaccharomyces pombe* [*mcherry-atb2::hph leu1-32 ura4-D18 h-*] | Dr. Mithilesh Mishra (DBS, TIFR) |
| CCDY375 | CCDY346/pREP42- HpFtsZ-G213A-GFP | This work |
| CCDY376 | CCDY346/pREP42- SaFtsZ-G196A-GFP | This work |
| CCDY404 | CCDY346/pREP42- HpFtsZ-T216E-GFP | This work |
| *Bacterial Strains* | | |
| CCD190 | DH10B; F – *mcrA* Δ(*mrr-hsdRMS-mcrBC*) φ80*lacZ*ΔM15 *ΔlacX74 recA1 endA1 araD139* Δ (*ara-leu*)7697 *galU galK* λ– *rpsL*(StrR) *nupG* | Invitrogen |
| CCD249 | *Staphylococcus aureus* ATCC 25923 | Lab Stock |
| -- | *Helicobacter pylori* 26695 | Dr. Asima Bhattacharyya |

| Plasmids | | |
|---|---|---|
| pCCD4 | pREP42- EcFtsZ-GFP | Srinivasan, et al. 2008 |
| pCCD51 | pREP42C- GFP | This work |
| pCCD712 | pREP42- HpFtsZ-GFP | This work |
| pCCD713 | pREP42- SaFtsZ-GFP | This work |
| pCCD716 | pREP42- SaFtsZ-G213A-GFP | This work |
| pCCD753 | pREP42- HpFtsZ-G196A-GFP | This work |
| pCCD838 | pREP42- HpFtsZ-T216E-GFP | This work |

**Supplementary Table – S2**

**List of oligonucleotides:**

| | | |
|---|---|---|
| RSO73 | 5' GCCTCCCCCGGGAACAACAACAACGCTAGCATGAGTAAAGGAGAAGAACTTTTC 3' | Forward primer for GFP (with SmaI site) used for creating pREP42C-GFP |
| RSO74 | 5' TTATTTGTATAGTTCATCCATGC 3' | Rev primer for GFP_STOP |
| RSO468 | 5' ATAGTCGCTTTGTTAAATCATATGGTTCATCAATCAGAG 3' | Forward primer having NdeI site for cloning of HpFtsZ |
| RSO469 | 5' AGCGTTGTTGTTGTTCCCGGGATCCTCATTGTCTTGCTGGATTC 3' | Reverse primer having BamHI site for cloning of HpFtsZ |
| RSO470 | 5' ATAGTCGCTTTGTTAAATCATATGTTAGAATTTGAACAAGGATTTAATC 3' | Forward primer having NdeI site for cloning of SaFtsZ |
| RSO471 | 5' AGCGTTGTTGTTGTTCCCGGGATCCTTATTACGTCTTGTTCTTCTTG 3' | Reverse primer having BamHI site for cloning of SaFtsZ |
| RSO650 | 5' GCTGTGAGTGCCATTTCTACTATCATCACTAAACC 3' | Forward primer for cloning of G213A_HpFtsZ |
| RSO651 | 5' GTAGAAATGGCACTCACAGCCCTAACCAAG 3' | Reverse primer for cloning of G213A_HpFtsZ |

| | | |
|---|---|---|
| RSO648 | 5' GTGTACAAGCTATCTCAGACTTAATCGCTGTTTC 3' | Forward primer for cloning of G196A_SaFtsZ |
| RSO649 | 5' CTGAGATAGCTTGTACACCTTGGCGTAAC 3' | Reverse primer for cloning of G196A_SaFtsZ |
| RSO808 | 5' GCATTTCTGAGATCATCACTAAACCCGG 3' | Forward primer for cloning of T216E_HpFtsZ |
| RSO809 | 5' GTGATGATCTCAGAAATGCCACTCACAG 3' | Reverse primer for cloning of T216E_HpFtsZ |

**Supplementary Table – S3**

**List of reagents and chemicals**

| Reagents | Manufacturer/ Catalog No. |
|---|---|
| LB medium | (Difco – 244620) |
| Brucella broth | BD 211088 |
| Fetal bovine serum | HiMedia – RM9970 |
| Carbenicillin | Sigma – C1389 or MP Biomedicals - 0219509201 |
| Kanamycin | Sigma – 60615 |
| Chloramphenicol | Sigma – C0857 |
| YES Agar | Formedium$^{TM}$ - PCM0410 |
| YES Broth | Formedium$^{TM}$ - PCM0310 |
| Edinburg minimal medium (EMM Agar or EMM Broth) | Formedium$^{TM}$ - PMD0210 |
| Adenine | Formedium$^{TM}$ - DOC0229 |
| Histidine | Formedium$^{TM}$ - DOC0144 |
| Leucine | Formedium$^{TM}$ - DOC0157 |
| Uracil | Formedium$^{TM}$ - DOC0214 |
| Thiamine | Sigma - T4625 |
| PC190723 | Merck - 344580 |
| Sanguinarine | Sigma - S5890 |
| Berberine | Sigma - B3251 |
| DMSO | Sigma - 317275 |
| Genomic DNA Isolation kit | HiMedia Labs - MB505 |
| Q5 Polymerase | NEB - M0492S |

| | |
|---|---|
| 96-well plate | Corning - CLS3370 |
